# Temporal landscape of mitochondrial proteostasis governed by the UPR^mt^

**DOI:** 10.1101/2022.11.30.518286

**Authors:** Louise Uoselis, Runa Lindblom, Marvin Skulsuppaisarn, Grace Khuu, Thanh N. Nguyen, Danielle L. Rudler, Aleksandra Filipovska, Ralf B. Schittenhelm, Michael Lazarou

**Affiliations:** Department of Biochemistry and Molecular Biology, Biomedicine Discovery Institute, Monash University, Melbourne, Australia; Walter and Eliza Hall Institute of Medical Research, Parkville, Victoria, Australia; Harry Perkins Institute of Medical Research and ARC Centre of Excellence in Synthetic Biology, Nedlands, Western Australia, Australia; Telethon Kids Institute, Northern Entrance, Perth Children’s Hospital, Nedlands, Western Australia, Australia; Monash Proteomics and Metabolomics Facility, Department of Biochemistry and Molecular Biology, Biomedicine Discovery Institute, Monash University, Melbourne, Australia.; Aligning Science Across Parkinson’s Collaborative Research Network, Chevy Chase, MD 20185, USA

## Abstract

Breakdown of mitochondrial proteostasis activates quality control pathways including the mitochondrial unfolded protein response (UPR^mt^) and PINK1/Parkin mitophagy. However, beyond the upregulation of chaperones and proteases, we have a limited understanding of how the UPR^mt^ remodels and restores damaged mito-proteomes. Here, we have developed a functional proteomics framework, termed MitoPQ (**Mito**chondrial **P**roteostasis **Q**uantification), to dissect the UPR^mt^’s role in maintaining proteostasis during stress. We discover essential roles for the UPR^mt^ in both protecting and repairing proteostasis, with oxidative phosphorylation metabolism being a central target of the UPR^mt^. Transcriptome analyses together with MitoPQ reveal that UPR^mt^ transcription factors drive independent signaling arms that act in concert to maintain proteostasis. Unidirectional interplay between the UPR^mt^ and PINK1/Parkin mitophagy was found to promote oxidative phosphorylation recovery when the UPR^mt^ failed. Collectively, this study defines the network of proteostasis mediated by the UPR^mt^ and highlights the value of functional proteomics in decoding stressed proteomes.

## INTRODUCTION

Mitochondrial quality control involves repair and removal processes that maintain mitochondrial health in the face of stress. Failure to maintain mitochondrial health has been implicated in the pathology of several human diseases ranging from neurodegenerative diseases, including Parkinson’s disease and Alzheimer’s disease to diabetes and cancer (Gaude and Frezza, 2016; Narendra *et al*., 2008; Silva *et al*., 2000; Wang *et al*., 2014). The fundamental biology of mitochondria, including metabolism and energy generation, is dictated by protein machineries whose proteostasis must be maintained to prevent dysfunction. The mitochondrial unfolded protein response (UPR^mt^) and PINK1/Parkin mitophagy are triggered in response to mitochondrial dysfunction but play opposing roles to restore proteostasis.

PINK1/Parkin mitophagy drives the degradation of severely damaged mitochondria through the *de novo* formation of autophagosomes that encapsulate damaged mitochondria before delivering them to lysosomes for degradation (Kane *et al*., 2014; Lazarou *et al*., 2015; Narendra *et al*., 2010; Nguyen *et al*., 2016). In contrast, the UPR^mt^ is a repair-driven process which induces the transcription of nuclear-encoded factors including chaperones and proteases to repair the protein folding environment of mitochondria (Zhao *et al*., 2002). While both the UPR^mt^ and PINK1/Parkin mitophagy are activated in response to protein folding stress within mitochondria (Fiesel *et al*., 2017; Jin and Youle, 2013), the interplay and crosstalk between these pathways is not well understood. 19 The UPR^mt^ has been best characterized in the model organism *C. elegans,* which has revealed critical insights into the organismal and cellular roles of the UPR^mt^ (Bar-Ziv *et al*., 2020; Naresh and Haynes, 2019). In *C. elegans*, the transcription factors ATFS-1 and DVE-1 are key signaling factors for the UPR^mt^ that play several roles to restore mitochondrial health including metabolic rewiring and driving the expression of proteostasis associated genes (Haynes *et al*., 2007; Nargund *et al*., 2015; Nargund *et al*., 2012; Tian *et al*., 2016). In human cells, the orthologue of ATFS-1 was identified to be ATF5 (Fiorese *et al*., 2016). However, beyond ATF5, there has been significant evolutionary divergence in UPR^mt^ signaling between *C. elegans* and humans including expansion of transcription factors and additional signaling mechanisms (Anderson and Haynes, 2020). During the UPR^mt^ in human cells, the mitochondrial protein DELE1 undergoes protein-folding stress induced cleavage that releases it to the cytosol where it signals the activation of the UPR^mt^ (Ahola *et al*., 2022; Fessler *et al*., 2020; Guo *et al*., 2020). An important component of DELE1 signaling is upregulated expression of key mammalian UPR^mt^ associated transcription factors including CHOP and ATF4 (Ahola *et al*., 2022; Fessler *et al*., 2020; Guo *et al*., 2020), of which CHOP is completely absent from the *C. elegans* genome (Schulz and Haynes, 2015). Collectively, these transcription factors drive a largely undefined UPR^mt^ program that involves proteases and chaperones to repair mitochondrial damage (Aldridge *et al*., 2007; Fiorese *et al*., 2016; Quirós *et al*., 2017; Zhao *et al*., 2002). The signaling hierarchy between each transcription factor, in addition to the relative importance of CHOP, ATF4 and ATF5 with respect to whether they act alone or in concert in driving the mammalian UPR^mt^ is unknown.

Analyses of proteostasis during the UPR^mt^ so far have relied on quantitative proteomics and while these approaches can accurately quantify protein levels and provide important insights (Münch and Harper, 2016; Quirós *et al*., 2017), they do not capture changes to the functional proteome that are independent of protein quantity. Functional changes include post translational modifications, protein-protein interactions, or most importantly with respect to disruptions in proteostasis, protein folding status. Given this, we have a limited understanding of vulnerabilities within the mito-proteome during protein folding stress and the role played by the UPR^mt^ in maintaining proteostasis.

Here, we have developed a functional proteomics framework that enables quantitative analysis of mitochondrial proteostasis. Termed **Mito**chondrial **P**roteostasis **Q**uantification (MitoPQ), the experimental pipeline was used to generate a temporal profile of mitochondrial proteostasis before, during, and after recovery from proteostasis stress. Combining MitoPQ with knockout lines of CHOP, ATF4 and ATF5, or all three, revealed that the UPR^mt^ functions during two distinct phases of proteostasis stress: a protection phase that limits damage, followed by a repair phase which restores proteostasis. Mitochondrial transcription and translation, and oxidative phosphorylation metabolism were identified as being highly vulnerable to protein folding stress, with complex I having a high reliance on the UPR^mt^ for protection and repair. Through transcriptomics and MitoPQ, we find that CHOP, ATF4 and ATF5 function in concert by driving independent arms of the UPR^mt^ program to non-redundantly protect and repair mitochondrial proteostasis. Assessment of the interplay between PINK1/Parkin mitophagy and the UPR^mt^ identified a unidirectional signaling relationship between the pathways, and a role for the UPR^mt^ in sustaining PINK1/Parkin mitophagy activity during the repair phase. We also identify PINK1/Parkin mitophagy as a secondary response to proteostasis stress that serves to address mitochondrial dysfunction when the UPR^mt^ fails or is overwhelmed. Overall, our work defines the role of the UPR^mt^ in protecting and repairing mitochondrial metabolism while providing a functional proteomics framework for the analysis of stressed proteomes.

## RESULTS

### MitoPQ, a quantitative proteomics framework for the analysis of mitochondrial proteostasis

The UPR^mt^ is thought to play a fundamental role in mitochondrial proteostasis (Moehle *et al*., 2019), yet despite this it remains unknown to what extent the UPR^mt^ program protects or repairs proteostasis during stress, while the individual roles of the key UPR^mt^ transcription factors CHOP, ATF4 and ATF5 have not been explored. To address these gaps, we established MitoPQ (**Mito**chondrial **P**roteostasis **Q**uantification), a quantitative proteomics framework designed to measure mitochondrial protein solubility. The workflow, summarized in Figure 1A, utilizes isolated mitochondria and involves extracting soluble and insoluble proteins based on their separation with 0.5% Triton X-100. Detergent soluble and insoluble protein fractions are then extracted using 5% SDS, sonication, and low pH buffers (<pH 3). To each fraction, a standardized amount of the bacterial protein Ag85A from *Mycobacterium tuberculosis* is added which enables a relative comparison of peptide intensities in soluble and insoluble mitochondrial fractions, and the calculation of a relative total amount of protein that is used to measure shifts in protein solubility as a percentage of the total fraction.

**Figure 1.**
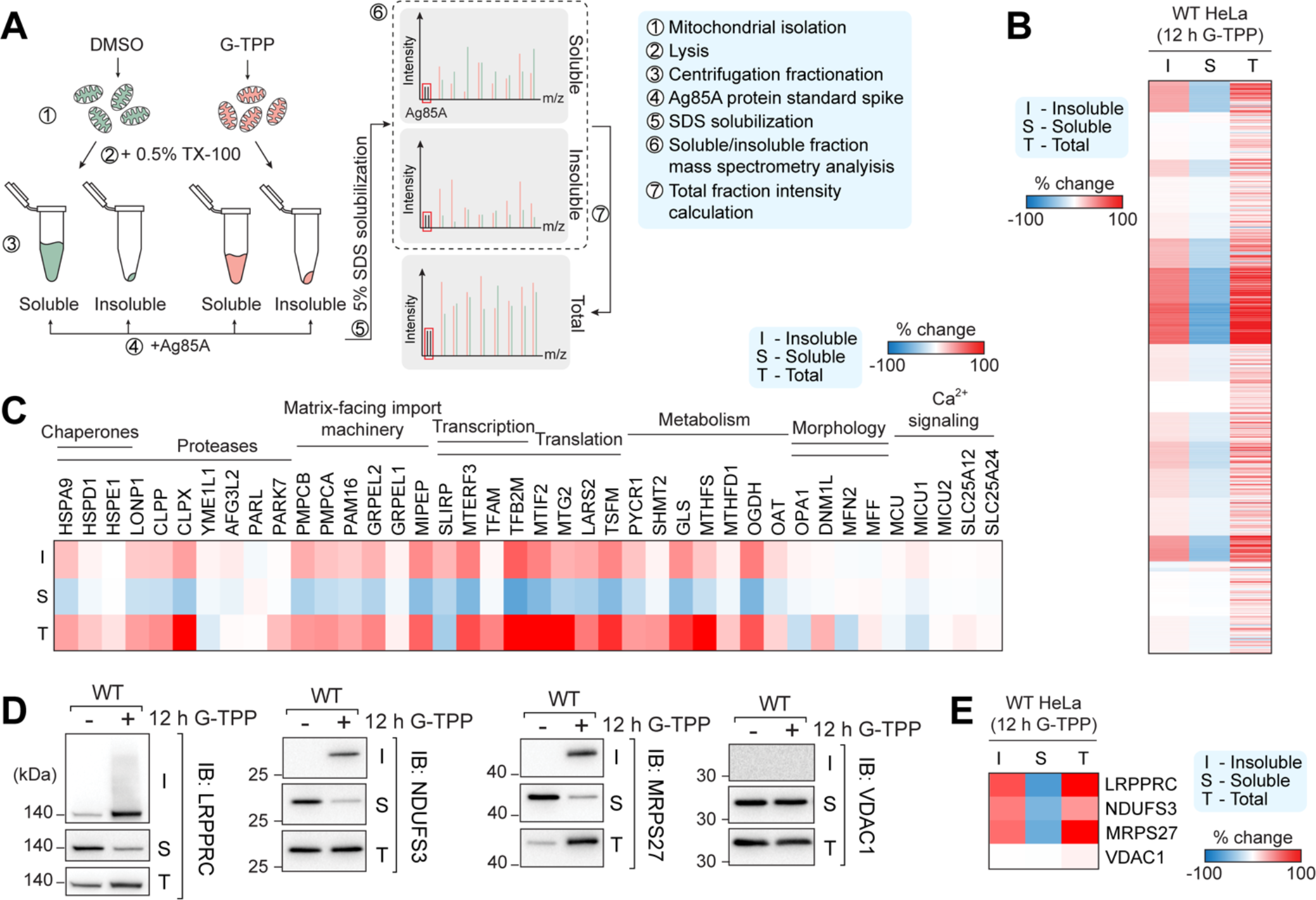
Developing an experimental pipeline to analyze endogenous mitochondrial proteostasis. (A) Workflow for proteostasis quantification of mitochondria. (B) Heat map analysis of mitochondrial protein solubility changes following 12 h G-TPP treatment. (C) Heat map of protein solubility changes grouped by mitochondrial process. (D) Immunoblot validation of solubility trends identified using MitoPQ analysis. (E) Heat map of solubility changes of proteins validated by immunoblot.

To validate the utility of MitoPQ in measuring mitochondrial proteostasis, MitoPQ analysis was performed on HeLa cells exposed to a mitochondrial protein folding stress. HeLa cells were treated for 12 h with G-TPP, a mitochondrial specific HSP90 inhibitor that induces mitochondrial protein misfolding (Fiesel *et al*., 2017; Münch and Harper, 2016). A total of 995 mitochondrial proteins were identified across the soluble and insoluble fractions (Figure 1B). A global clustered heat map analysis identified protein groups that displayed varied trends in changes to total protein levels and solubility following protein folding stress (Figure 1B). This included 221 mitochondrial proteins that became more insoluble following G-TPP treatment (>25% shift to the insoluble fraction), and 109 mitochondrial proteins whose solubility did not change but total protein levels did change, demonstrating varied changes to proteostasis across the mito-proteome (Figure 1B). Analysis of select mitochondrial machineries revealed that mitochondrial transcription and translation machineries, along with key metabolism associated factors including GLS, MTHFS, and OGDH, were acutely sensitive to solubility collapse following proteostasis stress (Figure 1C). In contrast, mitochondrial morphology and Ca^2+^ signaling processes were largely unaffected, indicating that sensitivity to proteostatic stress differs across the functional components of the mitoproteome (Figure 1C). To confirm the accuracy of MitoPQ, the solubility and total protein trends in cells after 12 h DMSO or 12h G-TPP treatment were analyzed by Western blotting (Figure 1D). Protein level trends in total, soluble, and insoluble fractions of select aggregating proteins (LRPPRC, NDUFS3, MRPS27) and a soluble control (VDAC1) assessed by immunoblotting (Figure 1D) mirrored those produced by MitoPQ analysis (Figure 1E), confirming the accuracy of the MitoPQ framework in quantifying changes to mitochondrial protein solubility.

### CHOP, ATF4, and ATF5 play essential roles to protect and repair mitochondrial proteostasis

Having established and validated MitoPQ, we next applied it to the fundamental question of how the UPR^mt^ maintains mitochondrial proteostasis through addressing the role played by UPR^mt^ transcription factors. HeLa cell KO lines of key UPR^mt^ transcription factors CHOP, ATF4, and ATF5, along with a triple KO (TKO) line lacking all three transcription factors were generated using CRISPR/Cas9 (ATF4 and ATF5), or TALEN (CHOP) mediated gene editing (Figure S1A and Table S1). First, we explored transcription factor induction in each KO line since previous reports have indicated that ATF4 is an upstream regulator of CHOP and ATF5 during the integrated stress response (ISR) (Ma *et al*., 2002; Teske *et al*., 2013). Immunoblot and mRNA analysis of transcription factor induction following G-TPP treatment of KO lines showed that in contrast to the canonical ISR in which ATF4 is the master regulator (Ma *et al*., 2002; Teske *et al*., 2013), each of CHOP, ATF4 and ATF5 were activated independently of each other during the UPR^mt^ (Figure 2A-C). ATF5 induction occurred post-translationally (Figure 2A- C), consistent with previous observations (Fiorese *et al*., 2016). These results demonstrate that CHOP, ATF4 and ATF5 are independently expressed during the UPR^mt^, and may therefore represent independent signaling arms of the UPR^mt^ program.

**Figure 2.**
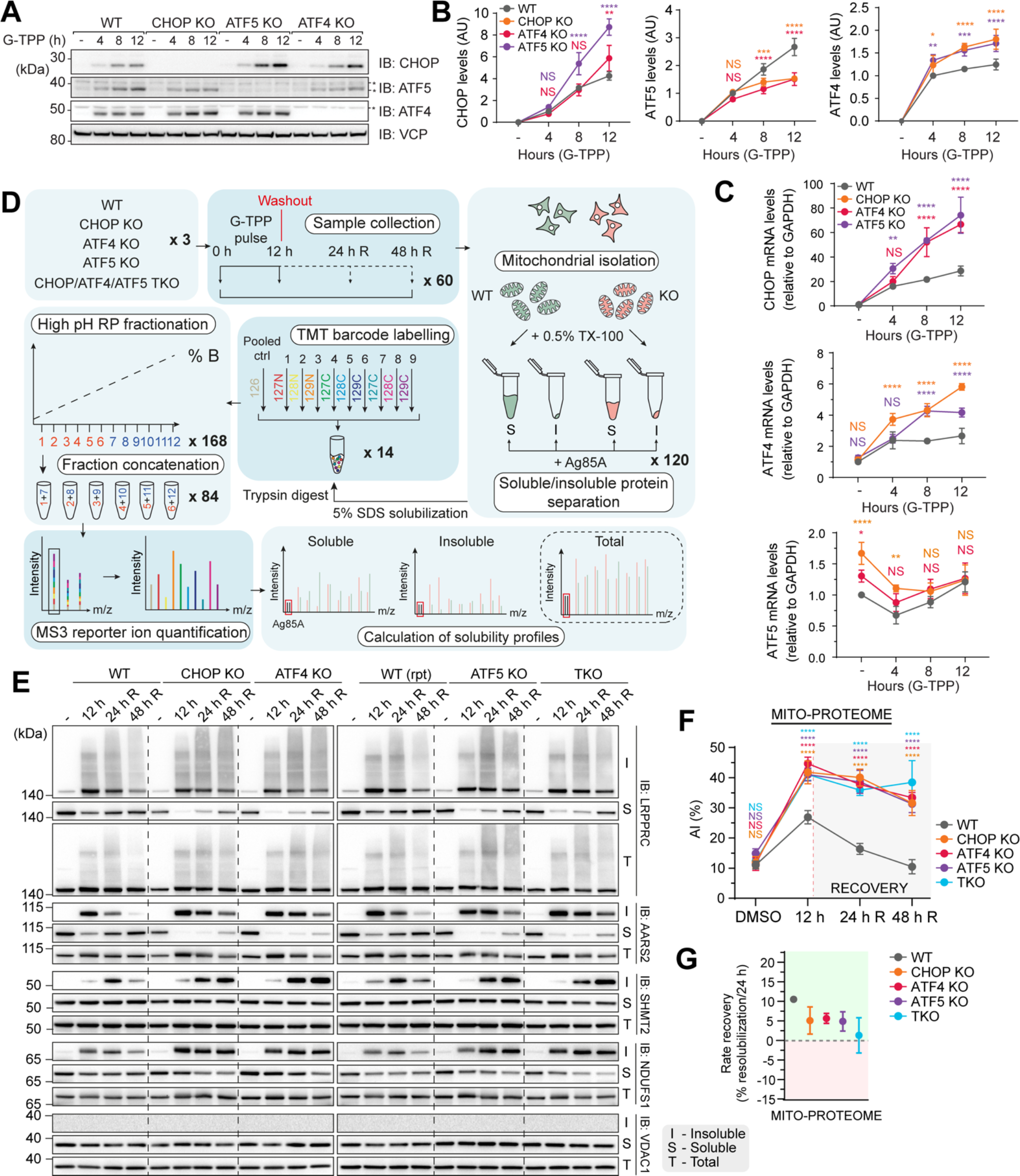
Independent signaling pathways driven by CHOP, ATF4 and ATF5 are required for UPR^mt^-driven proteostasis protection and repair. (A-C) WT, CHOP KO, ATF4 KO and ATF5 KO cells were treated with G-TPP and transcription factor expression was analyzed by immunoblot (IB; A) and (B) quantified. mRNA expression of each transcription factor was analyzed by qRT-PCR and quantified (C). (D) Experimental design. (E) IB validation of solubility trends identified in the experiment outlined in (D) using MitoPQ analysis. (F) Plot of mitoproteome aggregation indices (AIs) in WT, CHOP KO, ATF4 KO, ATF5 KO and TKO samples. (G) Plot of resolubilzation rates for WT, CHOP KO, ATF4 KO, ATF5 KO and TKO cells across the mitoproteome (linked to data in Table 1). Data in (B), (C), (F) and (G) are mean ± SD from three independent experiments. *p<0.05, **p<0.005. ***p<0.001, ****p<0.0001 and NS = not significant (two-way ANOVA).

**Table 1.** Statistical analysis of rate recovery trends across mitochondrial proteome and process groupings. Samples were analyzed relative to WT rate recovery per 24 h (‘Relative to WT’ column) or TKO rate recovery per 24 h (‘Relative to TKO’ column). ****p≤0.0001, ***p≤0.001, **p≤0.01, *p≤0.05, NS p>0.05 (one-way ANOVA). Data was generated from three independent experiments.

|  | Relative to WT |  |  |  | Relative to TKO |  |  |
| --- | --- | --- | --- | --- | --- | --- | --- |
|  | CHOP KO | ATF4 KO | ATF5 KO | TKO | CHOP KO | ATF4 KO | ATF5 KO |
| Amino acid metabolism | NS | NS | NS | ** | NS | NS | NS |
| Apoptosis | NS | NS | NS | NS | NS | NS | NS |
| Calcium Signaling and Transport | NS | NS | NS | NS | NS | NS | NS |
| Cardiolipin biosynthesis | NS | NS | ** | NS | NS | NS | * |
| Fatty Acid Biosynthesis & Elongation | NS | NS | * | *** | ** | ** | * |
| Fatty Acid Degradation & Beta-oxidation | NS | NS | NS | ** | NS | NS | NS |
| Fatty Acid Metabolism | * | ** | * | ** | NS | NS | NS |
| Fe-S Cluster Biosynthesis | NS | NS | NS | NS | NS | NS | NS |
| Folate & Pterin Metabolism | NS | NS | NS | NS | NS | NS | NS |
| Glycolysis | NS | NS | NS | * | NS | ** | NS |
| Heme Biosynthesis | NS | NS | NS | * | NS | NS | NS |
| Import & Sorting | ** | ** | * | ** | NS | NS | NS |
| Metabolism of Lipids & Lipoproteins | NS | * | * | * | NS | NS | NS |
| Metabolism of Vitamins & Co-Factors | NS | NS | NS | * | NS | NS | NS |
| Mitochondrial Carrier | NS | NS | NS | NS | NS | NS | NS |
| Mitochondrial Dynamics | NS | NS | NS | NS | NS | NS | NS |
| Mitochondrial Signaling | NS | NS | NS | NS | NS | NS | NS |
| Mitophagy | NS | NS | * | ** | NS | * | NS |
| Nucleotide Metabolism | NS | NS | NS | ** | NS | * | NS |
| Oxidative Phosphorylation | * | ** | * | *** | * | NS | * |
| Pentose Phosphate Pathway | NS | ** | NS | NS | NS | ** | NS |
| Protein Stability & Degradation | NS | NS | NS | NS | NS | NS | NS |
| Pyruvate Metabolism | NS | NS | NS | NS | NS | NS | NS |
| Replication & Transcription | ** | ** | ** | **** | ** | ** | ** |
| ROS Defence | NS | NS | NS | NS | NS | * | NS |
| Translation | *** | *** | *** | **** | * | * | ** |
| Transmembrane Transport | * | NS | NS | * | NS | NS | NS |
| TCA Cycle | NS | NS | NS | ** | NS | * | NS |
| Ubiquinone Biosynthesis | ** | *** | ** | **** | *** | ** | *** |
| Mito-proteome | * | NS | * | ** | NS | NS | NS |

We next addressed the contribution of each transcription factor to the protection and repair of mitochondrial proteostasis during stress. The MitoPQ framework was coupled with TMT multiplex barcode labelling and applied to transcription factor KO lines treated with G-TPP. Cells were analyzed at four different time points, including a 12 h DMSO treatment control, a 12 h G-TPP treatment to assess UPR^mt^ mediated protection against proteostasis damage, and 24 h and 48 h G-TPP wash out time points, termed recovery (R), to assess UPR^mt^-mediated proteostasis repair. A schematic summary of the experimental workflow is shown in Figure 2D. Temporal solubility profiles were calculated for 884 mitochondrial proteins across the entire dataset. Principle component analyses showed highly reproducible, time dependent proteostasis patterns across each cell line, with baseline DMSO samples clustering with recovery samples in WT cells, but not in KO lines (see ‘DMSO’ and 48 h R’, Figure S1B). Immunoblot analysis of select proteins identified by MitoPQ as highly aggregating (LRPPRC and AARS2), mildly aggregating (NDUFS1), or completely soluble (VDAC1), validated the temporal proteostasis profiles generated by MitoPQ analysis (Figure 2E).

Next, we compared the global proteostasis profiles across all KO cell lines and the WT control by calculating an aggregation index (AI) which represents the global average of protein aggregation in each cell line at each time point (Figure 2F). The aggregation index peaked in each cell line at 12 h G-TPP treatment (Figure 2F), with CHOP, ATF4, and ATF5 KOs displaying significantly higher levels of aggregation than the WT control (Figure 2F). Analysis of the recovery time points (24 h R and 48 h R) in WT cells showed that the aggregation index decreased at 24 h, and returned to the DMSO baseline by 48 h (Figure 2F), demonstrating an almost complete recovery from the protein folding stress. In contrast, each transcription factor KO cell line showed little reduction in the aggregation index at 24h recovery, and minimal reduction by 48 h (Figure 2F). The apparent lack of recovery in KO lines may be influenced by their high aggregation load at the 12 h treatment time point. To overcome this, we assessed global recovery rates as a percentage of aggregation reduction over a 24 h period (Figure 2G, Table 1) and found that all KO lines had an ∼50% or greater reduction in their rate of recovery relative to the WT control. These results demonstrate that the UPR^mt^ functions during two distinct phases of mitochondrial protein folding stress: 1) a protection phase that occurs during the insult to limit proteostasis damage, and 2) a repair phase that occurs following the removal of the insult to restore proteostasis.

We noted that the TKO line did not show an additive proteostasis defect, either during the protection or repair phases of the UPR^mt^ (Figure 2F-G). However, given that the aggregation index is a measure of average protein aggregation, we assessed whether TKO cells had a higher proportion of very highly aggregating proteins. TKO cells did not display an increased level of highly aggregating proteins which we classified as having >50% solubility shift (Figure S2). These results demonstrate that each of the CHOP, ATF4 and ATF5 signaling arms of the UPR^mt^ play equally important roles in protecting and repairing mitochondrial proteostasis.

### Mitochondrial processes are differentially affected by the loss of UPR^mt^ function

The global analysis in Figure 2F provided a broad overview of mitochondrial protein aggregation. However, we also wanted to gain information on specific mitochondrial processes to assess their sensitivity to aggregation in the presence/absence of a functional UPR^mt^ program. In addition, we wanted to address whether CHOP, ATF4, or ATF5 have any selectivity toward the protection or repair of specific mitochondrial processes. A broad heat map analysis was conducted on mitochondrial proteins grouped according to their functional processes using a list curated by (Kuznetsova *et al*., 2021) (Figure 3A). Following 12 h G-TPP treatment, the heat map revealed clusters of proteins with elevated aggregation in WT cells (Figure 3A), that were further aggregated in UPR^mt^ deficient transcription factor KO lines. The highly aggregating clusters displayed a resistance to repair in the absence of the UPR^mt^ (Figure 3A, 24 h and 48 h recovery). The heat map analysis indicates that certain mitochondrial processes are sensitive to protein folding stress and have a reliance on the UPR^mt^ for their proteostasis.

**Figure 3.**
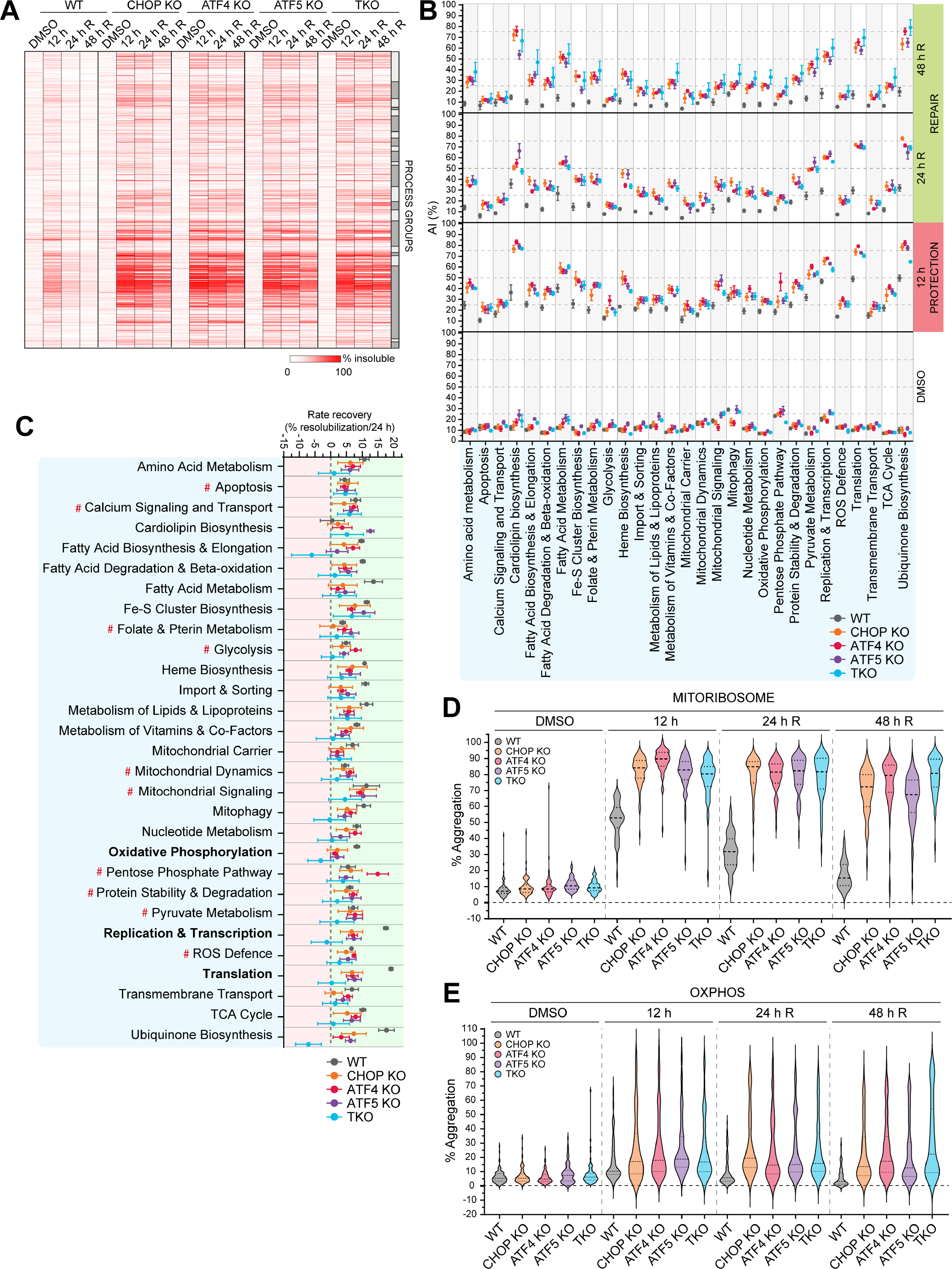
Each CHOP, ATF4 and ATF5 signaling pathway functions non-redundantly in driving UPR^mt^ mediated proteostasis protection and repair. (A) Heat map analysis clustered by mitochondrial process of solubility trends in WT, CHOP KO, ATF4 KO, ATF5 KO and TKO cells across the indicated timepoints. Alternating light and dark grey boxes correspond to individual process groups. (B) Aggregation indices (AI) of WT, CHOP KO, ATF4 KO, ATF5 KO and TKO cells were calculated and graphed for each mitochondrial process group (linked to data in Table 2). (C) Resolubilization rates for WT, CHOP KO, ATF4 KO, ATF5 KO and TKO cells were calculated and graphed for each mitochondrial process group. (D-E) Mean % aggregation of proteins belonging to the mitochondrial ribosome (D) or the OXPHOS machinery (E) were graphed by violin plot. Solid lines = median, dotted lines = quartiles. Data in (A), (D) and (E) represent mean data calculated from three independent experiments. Data in (B, C) are mean ± SD from three independent experiments.

**Table 2.** Statistical analysis of AI trends across mitochondrial proteome and process groupings. Samples were analyzed relative to WT aggregation index (AI) levels at each time point. ****p≤0.0001, ***p≤0.001, **p≤0.01, *p≤0.05, NS p>0.05 (two-way ANOVA). Data was generated from three independent experiments.

|  | DMSO |  |  |  | 12 h |  |  |  | 24 h recovery |  |  |  | 48 h recovery |  |  |  |
| --- | --- | --- | --- | --- | --- | --- | --- | --- | --- | --- | --- | --- | --- | --- | --- | --- |
|  | CHOP KO | ATF4 KO | ATF5 KO | TKO | CHOP KO | ATF4 KO | ATF5 KO | TKO | CHOP KO | ATF4 KO | ATF5 KO | TKO | CHOP KO | ATF4 KO | ATF5 KO | TKO |
| Amino acid metabolism | NS | NS | NS | NS | **** | **** | **** | **** | **** | **** | **** | **** | **** | **** | **** | **** |
| Apoptosis | NS | NS | NS | NS | **** | *** | *** | **** | **** | **** | *** | *** | NS | NS | NS | NS |
| Calcium Signaling and Transport | NS | NS | * | NS | ** | **** | ** | *** | **** | **** | **** | **** | NS | NS | NS | NS |
| Cardiolipin biosynthesis | NS | ** | **** | *** | **** | **** | **** | **** | **** | **** | **** | **** | **** | **** | **** | **** |
| Fatty Acid Biosynthesis & Elongation | NS | NS | **** | ** | **** | **** | **** | *** | **** | **** | **** | **** | **** | **** | **** | **** |
| Fatty Acid Degradation & Beta-oxidation | NS | NS | NS | NS | **** | **** | **** | **** | **** | **** | **** | **** | **** | **** | **** | **** |
| Fatty Acid Metabolism | NS | * | **** | *** | **** | **** | **** | **** | **** | **** | **** | **** | **** | **** | **** | **** |
| Fe-S Cluster Biosynthesis | NS | * | NS | * | **** | **** | **** | **** | **** | **** | **** | **** | **** | **** | *** | **** |
| Folate & Pterin Metabolism | NS | NS | **** | * | **** | **** | **** | **** | **** | **** | **** | **** | **** | **** | **** | **** |
| Glycolysis | NS | NS | ** | NS | * | **** | ** | * | *** | ** | ** | ** | NS | NS | NS | ** |
| Heme Biosynthesis | NS | NS | ** | NS | **** | **** | **** | **** | **** | **** | **** | **** | **** | **** | **** | **** |
| Import & Sorting | NS | NS | * | NS | NS | ** | ** | *** | **** | **** | **** | **** | *** | **** | ** | **** |
| Metabolism of Lipids & Lipoproteins | NS | NS | **** | ** | *** | *** | **** | *** | **** | **** | **** | **** | ** | *** | *** | *** |
| Metabolism of Vitamins & Co-Factors | NS | NS | * | NS | **** | **** | **** | **** | **** | **** | **** | **** | **** | **** | **** | **** |
| Mitochondrial Carrier | NS | NS | NS | NS | *** | **** | *** | ** | **** | **** | *** | **** | ** | **** | *** | ** |
| Mitochondrial Dynamics | NS | NS | NS | NS | *** | **** | **** | *** | **** | * | **** | *** | * | NS | NS | ** |
| Mitochondrial Signaling | NS | ** | ** | *** | **** | **** | **** | **** | **** | **** | **** | **** | *** | *** | **** | **** |
| Mitophagy | **** | ***<br>* | NS | NS | NS | * | NS | NS | **** | ** | **** | **** | * | NS | ** | **** |
| Nucleotide Metabolism | NS | NS | NS | NS | **** | **** | **** | **** | **** | **** | **** | **** | **** | **** | **** | **** |
| Oxidative Phosphorylation | NS | NS | NS | NS | *** | *** | *** | ** | **** | **** | **** | **** | **** | **** | **** | **** |
| Pentose Phosphate Pathway | NS | NS | * | ** | *** | **** | *** | **** | ** | **** | **** | * | NS | * | ** | **** |
| Protein Stability & Degradation | NS | NS | ** | NS | **** | **** | **** | **** | **** | **** | **** | **** | **** | **** | **** | **** |
| Pyruvate Metabolism | ** | NS | ** | NS | **** | **** | **** | **** | **** | **** | **** | **** | **** | **** | **** | **** |
| Replication & Transcription | NS | NS | ** | * | **** | **** | **** | **** | **** | **** | **** | **** | **** | **** | **** | **** |
| ROS Defence | NS | NS | NS | NS | **** | **** | **** | **** | **** | **** | **** | **** | ** | ** | * | **** |
| Translation | NS | NS | NS | NS | **** | **** | **** | **** | **** | **** | **** | **** | **** | **** | **** | **** |
| Transmembrane Transport | NS | NS | NS | NS | NS | *** | ** | ** | **** | NS | *** | **** | ** | NS | * | *** |
| TCA Cycle | NS | NS | NS | NS | **** | **** | **** | **** | **** | **** | **** | **** | **** | **** | **** | **** |
| Ubiquinone Biosynthesis | NS | NS | NS | NS | **** | **** | **** | **** | **** | **** | **** | **** | **** | **** | **** | **** |
| Mito-proteome | NS | NS | * | NS | ** | *** | ** | ** | *** | *** | ** | *** | ** | *** | *** | * |

To clearly identify and characterize the mitochondrial processes that belonged to the aggregation sensitive protein clusters in Figure 3A, an aggregation index was calculated for each functional process group and plotted across the protection and recovery phases (Figure 3B, Table 2). In WT cells, all mitochondrial process groups displayed an elevated aggregation index following 12 h G-TPP treatment except for *Apoptosis*, *Glycolysis*, *Mitochondrial Carrier* and *Transmembrane Transport* (Figure 3B). High aggregation index values (>40%) at 12 h G-TPP treatment were observed for metabolism related processes including, *Fatty Acid Metabolism* and *Ubiquinone Biosynthesis*, in addition to *Replication and Transcription*, and *Translation* (Figure 3B), indicating high sensitivity of these mitochondrial processes to proteostasis stress. In transcription factor KO lines, all mitochondrial process groups had aggregation indexes that were above the WT control to varying degrees, with metabolism related processes such as *Fe-S biosynthesis*, *Cardiolipin Biosynthesis*, *Ubiquinone Biosynthesis,* and *Translation* showing a very high reliance on the UPR^mt^ during the protection phase (Figure 3B).

Analysis of the 24 h and 48 h recovery time points, representing the proteostasis repair phase of the UPR^mt^, showed that WT cells returned to baseline aggregation indexes by 48 h recovery across all process groups (Figure 3B). In contrast, with the exception of *Apoptosis*, and *Calcium Signaling and Transport*, aggregation indexes remained high in all transcription factor KO lines demonstrating widespread defects in proteostasis repair. Recovery rates were also calculated for each mitochondrial process group to further assess UPR^mt^ mediated proteostasis repair (Figure 3C, Table 1). Unexpectedly, this analysis revealed that multiple process groups in transcription factor KO cells had similar recovery rates to WT, including *Folate and Pterin Metabolism* and *Glycolysis* (see process groups marked by # in Figure 3C), demonstrating a reliance on the UPR^mt^ for protection (Figure 3B) but not repair. In contrast, several mitochondrial processes required the UPR^mt^ for both protection and repair, and these included *Oxidative Phosphorylation*, *Replication and Transcription*, and *Translation* process groupings (Figure 3C, highlighted in bold text). The TKO line had broadly similar protection and repair defects to the single KO lines (Figure 3A-C, Tables 1-2), although decreased recovery rates were observed for *Fatty Acid Metabolism*, *Ubiquinone Biosynthesis, Replication and Transcription, and Translation* (Figure 3C), indicating partial redundancy of the transcription factors for these mitochondrial processes. Overall, CHOP, ATF4, or ATF5 did not show a preference in protecting or repairing specific mitochondrial processes, with all three transcription factors playing equally important roles in maintaining proteostasis. In addition, we find a divergence in mitochondrial processes in terms of their requirement for the CHOP, ATF4 and ATF5 driven UPR^mt^ program, with some requiring it solely for protection from stress (e.g. glycolysis), while the majority were reliant on the UPR^mt^ for both protection and repair.

### The UPR^mt^ protects and repairs complex I to maintain OXPHOS activity

Oxidative Phosphorylation (OXPHOS) and translation were major mitochondrial process groups that were found to rely on the UPR^mt^ for their protection and repair (Figure 3). These fundamental mitochondrial processes consist of protein complexes that drive their activity. For example, the *Oxidative Phosphorylation* process group consists of multiple machineries of the electron transport chain (complexes I-IV) in addition to the ATP synthase complex (complex V). We asked whether all components of the translation machinery and all complexes of oxidative phosphorylation machineries were equally disrupted by protein folding stress, and to what degree the UPR^mt^ provides protection and repair. The mitoribosome (94% of subunits identified by MitoPQ) displayed broad aggregation across all protein components belonging to both the 39S large subunit and the 28S small subunit (Figure 3D). All proteins belonging to both the 39S and 28S subunits of the mitoribosome underwent proteostasis repair in WT cells but not in UPR^mt^ defective KO lines (Figure 3D), which was consistent with their impaired recovery rates (Figure 3C). In contrast to the mitoribosome (Figure 3D), aggregation analysis of the entire OXPHOS machinery (74% of subunits identified by MitoPQ) showed only a moderate median level of aggregation, but a large upper tail of highly aggregating proteins was observed (Figure 3E), revealing that a specific subset of OXPHOS proteins are highly sensitive to aggregation. To identify the highly aggregating OXPHOS proteins, violin plots were separated out by individual OXPHOS complexes (complexes I – V) (Figure 4A, B, and Figure S3A-C). This analysis revealed that in both WT and transcription factor KO cells, the highly aggregating OXPHOS proteins were primarily complex I subunits (compare Figure 4A to 4B and to Figure S3A-C). In UPR^mt^ defective cells, further aggregation of complex I subunits was observed relative to WT controls (Figure 4A), along with additional aggregation of CII–CV subunits that were otherwise not strongly aggregated in WT cells (Figure 4B and Figure S3A-C). Collectively, the KO lines displayed a failure to both protect and repair complex I proteostasis (Figure 4A), with the strongest defect observed during the repair phase of the UPR^mt^. Overall, this analysis has identified complex I as a vulnerable component of the OXPHOS machinery that is highly reliant on the UPR^mt^ for its protection and repair during stress. Combined with the observation that the UPR^mt^ also strongly protects and repairs processes that support OXPHOS (cardiolipin biosynthesis, fatty acid metabolism, ubiquinone biosynthesis and transcription/translation (Figure 3)), we conclude that the UPR^mt^ plays a fundamental role in maintaining OXPHOS metabolism during proteostasis stress.

**Figure 4.**
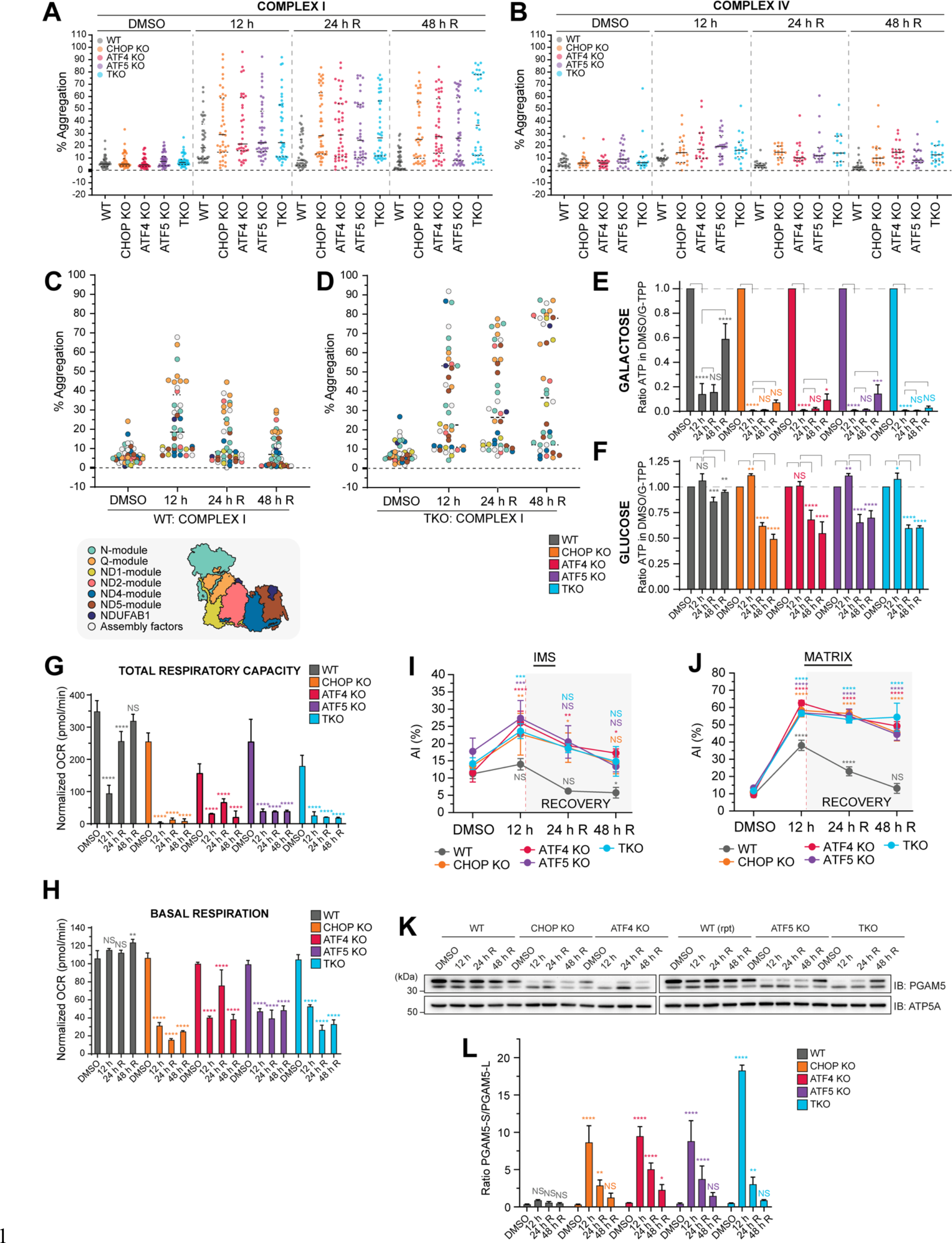
The UPR^mt^ protects and repairs OXPHOS solubility and activity during proteostasis stress. (A, B) Violin plots of the mean % aggregation of proteins comprising complex I (A) and complex IV (B) in WT, CHOP KO, ATF4 KO, ATF5 KO and TKO cells at the indicated timepoints. (C, D) Violin plots of the mean % aggregation of proteins comprising complex I, labelled according to complex I sub- module localization in WT (C) and TKO (D) cells at the indicated timepoints. (E, F) Ratios of cellular ATP levels in WT, CHOP KO, ATF4 KO, ATF5 KO and TKO cells treated in either galactose (E) or glucose (F) based media were calculated relative to DMSO levels at the indicated timepoints and graphed. (G, H) Oxygen consumption rates (OCR; pmol/min) of mitochondria isolated from WT, CHOP KO, ATF5 KO, ATF4 KO and TKO samples collected at the indicated timepoints were calculated using a Seahorse (Agilent) analyzer. OCR values of untreated cells were analyzed at each time point and used to normalize OCR values collected at each day of the experimental time course. Normalized OCR values were graphed and the total respiratory capacity (G) and basal respiration (H) were calculated and graphed for each sample. (I, J) Aggregation indices (AI) of intermembrane space (IMS; I) or matrix (J) localized proteins in WT, CHOP KO, ATF4 KO, ATF5 KO and TKO cells were calculated and graphed. (K, L) Uncleaved and cleaved PGAM5 levels were analyzed by immunoblot of mitochondria (K) isolated from WT, CHOP KO, ATF4 KO, ATF5 KO and TKO cells at the indicated time points. Ratios of cleaved/uncleaved PGAM5 were calculated within each sample and graphed (L). Data in (A – D) represent mean data calculated from three independent experiments. Data in (E – J) and (L) represent mean ± SD from three independent experiments. *p<0.05, **p<0.005. ***p<0.001, ****p<0.0001 and NS = not significant (two-way ANOVA, relative to DMSO control (G-J, L)).

We noted that complex I subunit aggregation did not display the same whole-complex aggregation trend observed with the mitoribosome (compare Figure 3E with Figure 3D). Given that complex I consists of distinct assembly modules (Formosa *et al*., 2018), the module locations of aggregating subunits were mapped onto violin plots to provide a spatial understanding of the aggregation sensitive areas of complex I (Figure 4C-D, Figure S3D-F). In WT cells, subunits with elevated aggregation were localized primarily in the matrix-facing N- and Q-modules involved in electron transfer (Figure 4C). Interestingly, transcription factor KO lines showed elevated aggregation of additional membrane-bound complex I subunits belonging to the ND-1 and ND-5 modules involved in proton pumping (Wirth *et al*., 2016), demonstrating a more widespread collapse of complex I proteostasis in the absence of each signaling arm of the UPR^mt^ (Figure 4D, Figure S3D-F).

To investigate the functional importance of UPR^mt^ mediated protection and repair of OXPHOS proteostasis, and simultaneously assess the accuracy of MitoPQ in determining the functional mito- proteome, we analyzed OXPHOS metabolic activity. First, ATP levels were measured in cells grown in galactose media which drives OXPHOS dependent ATP production (Figure 4E). Relative to the WT control, all UPR^mt^ defective KO lines had almost undetectable levels of ATP during protein folding stress (Figure 4E), they also failed to restore ATP levels following 24 h and 48 h recovery. Indeed, such was the severity of mitochondrial dysfunction, that KO lines cultured in glucose also displayed significantly reduced levels of ATP during the recovery phase of G-TPP treatment (Figure 4F). To directly measure OXPHOS activity, complex I stimulated oxygen consumption rates were analyzed using isolated mitochondria (Figure 4G-H and Figure S3G). In WT cells, there was a robust recovery of total respiratory capacity within 24 h of stress removal (Figure 4G), while basal respiration remained undisrupted across all time-points (Figure 4H). In contrast, all UPR^mt^ defective KO lines had severely defective respiratory capacity that was unrecoverable (Figure 4G), highlighting the importance of each of CHOP, ATF4 and ATF5 UPR^mt^ arms. Overall, these results reveal the fundamentally important role that the UPR^mt^ plays in maintaining OXPHOS metabolic activity during protein folding stress by protecting and repairing its proteostasis. The results also highlight the utility of the MitoPQ framework in measuring the functional mito-proteome.

### The UPR^mt^ is dispensable for IMS proteostasis repair

Inhibition of mitochondrial HSP90 using G-TPP provides a protein folding stress that originates in the mitochondrial matrix. We hypothesized that failure to contain a protein folding stress originating from the matrix, such as when the UPR^mt^ is defective or overwhelmed, can result in the stress spreading to the inter membrane space (IMS) compartment of mitochondria. To test this hypothesis, the levels of protein aggregation in the matrix and IMS compartments of mitochondria were compared between WT controls and UPR^mt^ defective KOs (Figure 4I-J). This analysis showed that indeed, when the UPR^mt^ is defective, a protein folding stress originating in the matrix can spread to the IMS compartment (Figure 4I). However, in contrast to the matrix compartment in which UPR^mt^ deficient cells showed little to no evidence of repair (Figure 4J), the IMS compartment showed recovery that reached close to the DMSO treated control in UPR^mt^ deficient cells (Figure 4I).

The cleavage and release of DELE1 from the IMS by the stress-activated protease OMA1 was identified as a key step of UPR^mt^ signaling (Fessler *et al*., 2020; Guo *et al*., 2020). OMA1 also cleaves the serine/threonine phosphatase PGAM5 (Sekine *et al*., 2012; Wai *et al*., 2016). Therefore, as an alternative readout of stress within the IMS, we assessed the ratio of inner membrane localized PGAM5-long (PGAM5-L) and cleaved PGAM5-short (PGAM5-S). Minimal IMS based stress was observed in WT cells based on PGAM5 cleavage analysis (Figure 4K-L). In contrast, all KO lines displayed significant levels of stress (Figure 4K-L). A further significant increase in IMS stress was observed in TKO cells relative to single transcription factor KOs (Figure 4K-L), indicating some redundancy in the roles of each CHOP, ATF4 and ATF5 in IMS protection. Interestingly, there was a robust recovery of the PGAM5- L/PGAM5-S ratio in all KO lines (Figure 4K-L). The UPR^mt^ therefore plays an important role to protect the IMS from proteostasis stress originating in the matrix, but is dispensable for recovery from this stress.

### CHOP, ATF4, and ATF5 govern common processes via both unique and overlapping genetic programs

Given that the loss of either CHOP, ATF4, and ATF5 resulted in near identical proteostasis defects (Figures 2-4), we asked whether each transcription factor functioned via identical or independent transcriptional programs. To address this, we conducted transcriptome analyses of cells treated with G- TPP for 12 h. In WT cells, the overall UPR^mt^ program significantly altered the expression 4610 genes (∼27% of the detected transcriptome), with similar numbers of upregulated and downregulated genes (Figure 5A), but the highest magnitude of change was observed for upregulated genes (Figure 5B). Upregulated genes were associated with GO processes linked to protein refolding, response to unfolded protein, and histone demethylase activity (Figure 5C, red), while downregulated genes were linked to histone methylation, transcription, and cell cycle (Fig 5C, blue). Pathway analysis of the transcriptome using KEGG identified cell cycle, ubiquitin mediated proteolysis, and Wnt signaling, among others, as UPR^mt^ regulated pathways (Figure 5D). Wnt signaling has been linked to cell-to-cell communication of the UPR^mt^ in *C. elegans* (Zhang *et al*., 2018), and may represent an evolutionary conserved node of the UPR^mt^ program. A more detailed analysis of Wnt signaling pathways revealed diverse gene expression patterns of upregulated and downregulated factors (Figure S4A), with the Wnt/Ca^2+^ signaling pathway showing an overall upregulation. Given the precedence of Wnt signaling in *C. elegans*, the link between the mammalian UPR^mt^ and Wnt is an interesting area for future exploration. To address whether the transcriptional changes resulted directly from UPR^mt^ signaling, or from pleiotropic signals arising from mitochondrial dysfunction, we analyzed the transcriptome of UPR^mt^ signaling deficient DELE1 KO cells (Fessler *et al*., 2020; Guo *et al*., 2020). An almost complete abolition of the UPR^mt^ transcriptional program was observed in DELE1 KOs (Figure S4B-C), confirming that the cellular pathways and processes identified above were a direct consequence of UPR^mt^ signaling originating from mitochondria.

**Figure 5.**
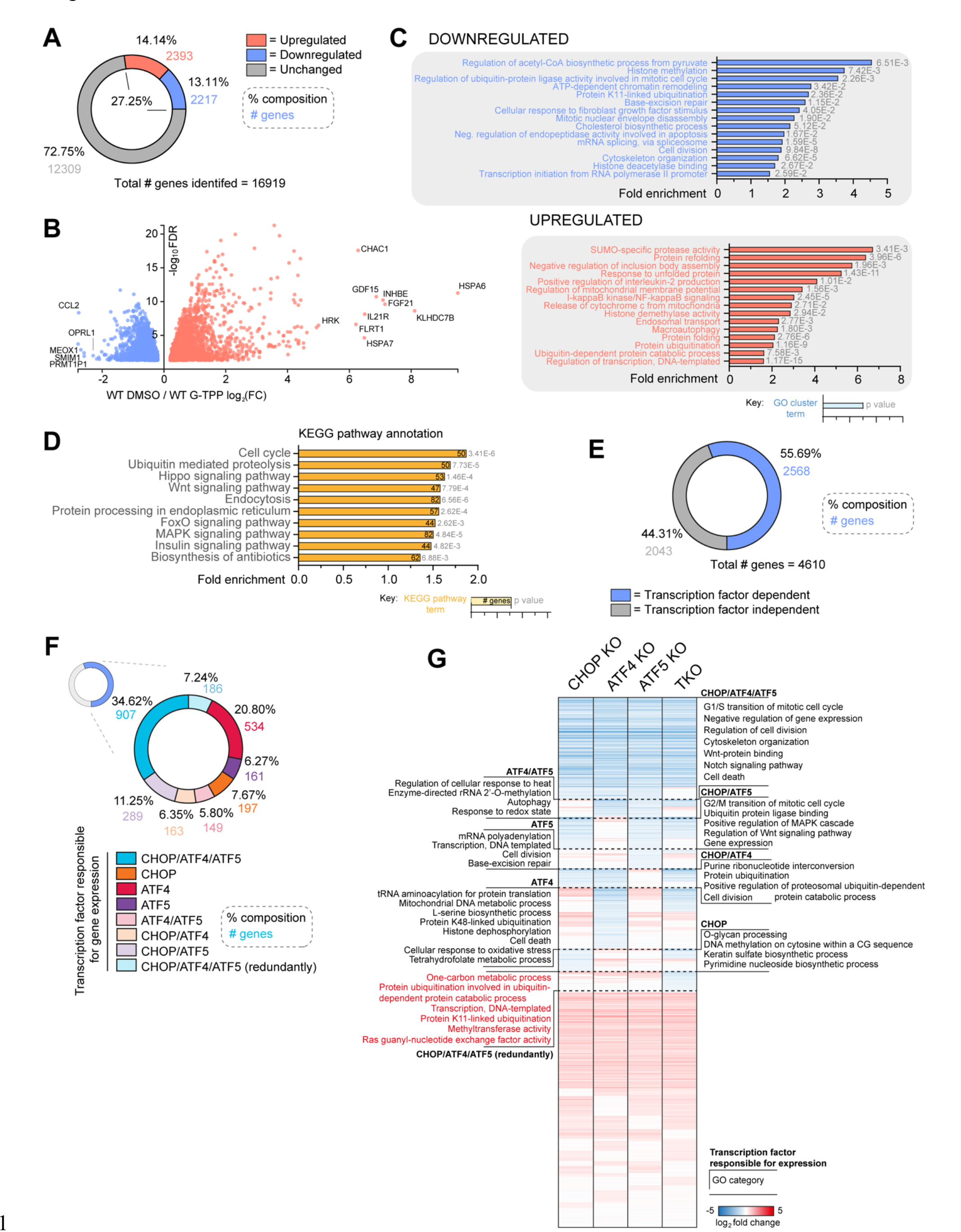
CHOP, ATF4 and ATF5 exert a mosaic pattern of regulatory genetic control to drive the UPR^mt^. (A) Pie chart breakdown of the percentage and number of genes in the transcriptome significantly altered or unchanged with UPR^mt^ induction. (B) Genes significantly upregulated or downregulated across the transcriptome in WT cells treated with G-TPP for 12 h relative to WT cells treated with DMSO for 12 h were defined as the WT UPR^mt^ transcriptome and graphed. The top 10 upregulated and top 5 downregulated genes in the WT UPR^mt^ transcriptome are labelled in (B). (C) Clustered gene ontology (GO) analysis of gene groups upregulated or downregulated in expression in the UPR^mt^ transcriptome. Gene clusters have been labelled using the most significantly enriched individual GO category in each cluster group. (D) KEGG pathway analysis of WT UPR^mt^ transcriptome trends. (E) Pie chart breakdown of UPR^mt^ transcriptome genes that were unchanged (‘Transcription factor independent’) or significantly decreased in expression in either CHOP KO, ATF4 KO, ATF5 KO or TKO cells treated with G-TPP for 12 h (‘Transcription factor dependent’). (F) Compositional breakdown displayed by pie chart of the overlapping regulatory patterns of CHOP, ATF4 and ATF5 in the transcription factor dependent portion of the UPR^mt^ transcriptome. (G) Heat map analysis of gene expression trends across the UPR^mt^ transcriptome in CHOP KO, ATF4 KO, ATF5 KO and TKO cells treated with G-TPP for 12 h, calculated relative to gene expression trends of WT cells treated with G-TPP for 12 h. Significantly enriched GO categories for each regulatory subgroup are labelled along the corresponding section of the heat map. Data in (A – G) represent mean data calculated from three independent experiments. GO analysis in (C) and (G) was performed using the Biological Process and Molecular Function subcategories in DAVID v6.8 (Huang *et al*., 2009). KEGG pathway analysis in (D) was performed using ShinyGO V0.75 (Ge *et al*., 2019).

We next analyzed the transcriptome of transcription factor KO lines to understand their contribution to the UPR^mt^. Of the 4610 significantly altered UPR^mt^ genes observed in WT cells, ∼56% were under the regulatory control of CHOP, ATF4 or ATF5 (Figure 5E). The 2568 genes regulated by CHOP, ATF4, and ATF5 displayed a mosaic pattern of regulation requiring either one, two, or all three transcription factors (Figure 5F-G), with a small subset of genes being regulated redundantly by the three transcription factors (Figure 5G, highlighted red). A large set of genes (907) required all three transcription factors (Figure 5F), followed by ATF4 alone (534), or a combination of CHOP and ATF5 (289). Of the three transcription factors, CHOP and ATF5 were the most related in terms of their UPR^mt^ signaling arms (Figure 5G, Figure S5A-B), whereas ATF4 had a more distinct UPR^mt^ signaling arm. Unique genetic footprints were also observed for each transcription factor (Figure 5G, see dashed boxed regions), but the uniquely regulated genes were associated with GO categories common to all three transcription factors including regulation of cell cycle, response to oxidative stress, ubiquitin signaling and proteasomal degradation, and Wnt signaling. Some unique GO categories were also detected, including tRNA aminoacylation for ATF4, mRNA polyadenylation for ATF5, and O-glycan processing for CHOP (Figure 5G). Genes associated with one carbon metabolism, which is remodeled in response to mitochondrial stress (Khan *et al*., 2017), were redundantly regulated by either CHOP, ATF4, and ATF5.

The global transcriptome analysis in Figure 5 can mask mitochondria associated genes since they represent only a small fraction of the total gene pool. To identify UPR^mt^-mediated mitochondrial changes that may have been masked in global gene analyses, a separate ontology analysis was performed on a curated gene set focused on mitochondria (Kuznetsova *et al*., 2021). Approximately 28% (370 of 1321) of the mitochondria associated gene set was altered in expression following UPR^mt^ activation (Figure 6A-B), and included upregulation of genes related to protein stability and degradation, protein import/sorting, and metabolism of amino acids, fatty acids, folate, vitamins and nitrogen (Figure 6C). In contrast, genes involved in OXPHOS, the TCA cycle and transcription/translation related processes were largely downregulated by the UPR^mt^ (Figure 6C).

**Figure 6.**
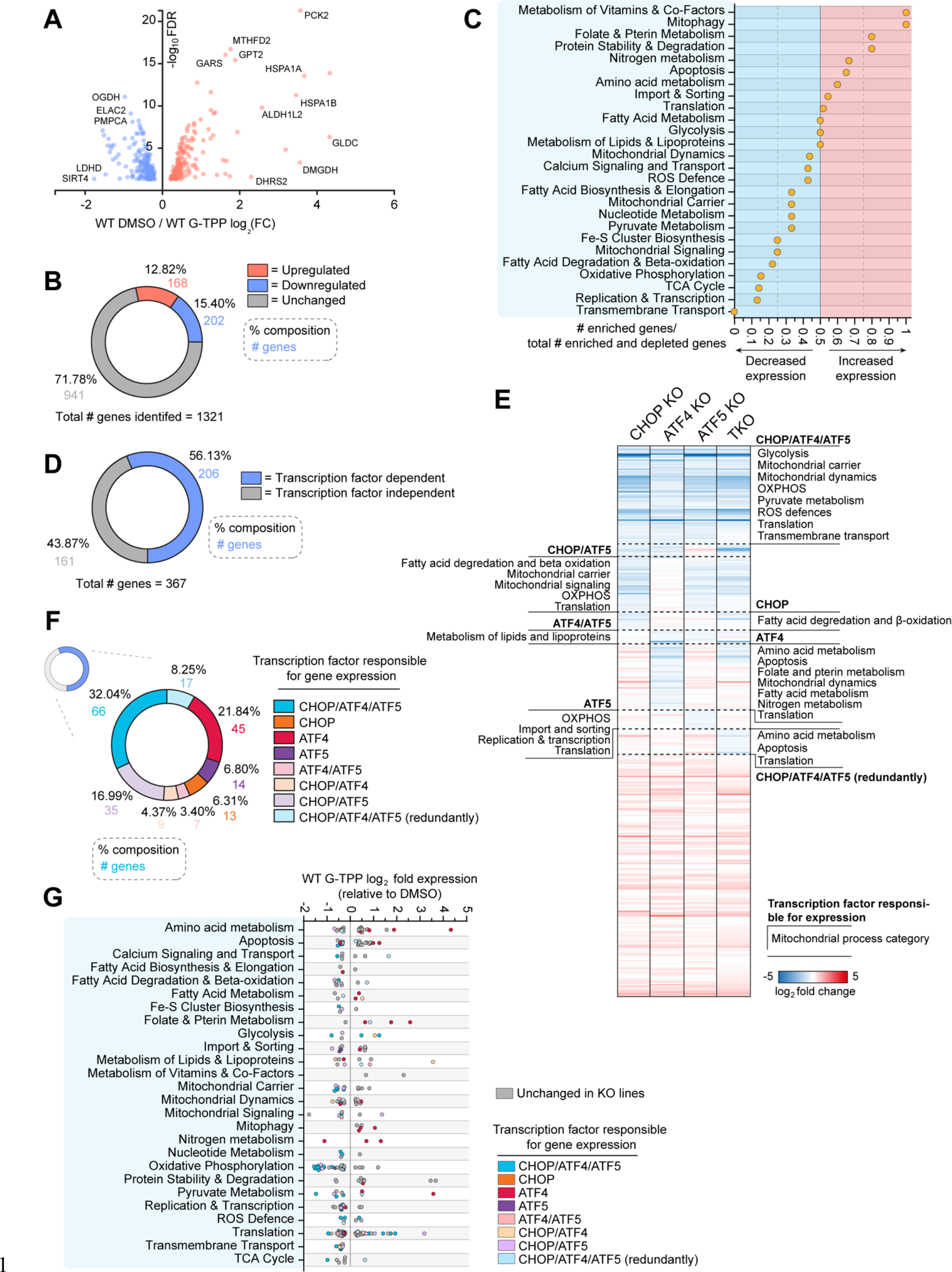
CHOP, ATF4 and ATF5 regulate mitochondrial gene expression through the UPR^mt^. (A) Graph of the expression changes of genes encoding mitochondrial proteins significantly altered in expression within the WT UPR^mt^ transcriptome. (B) Pie chart breakdown of the percentage and number of mitochondrial genes identified in the WT transcriptome that are altered or unchanged with UPR^mt^ induction. (C) Mitochondrial genes significantly altered in expression with UPR^mt^ induction were grouped by mitochondrial process and the relative number of genes in each process category showing increased expression with UPR^mt^ induction were graphed. (D) Pie chart breakdown of mitochondrial UPR^mt^ transcriptome genes that were unchanged (‘Transcription factor independent’) or significantly decreased in expression in either CHOP KO, ATF4 KO, ATF5 KO or TKO cells treated with G-TPP for 12 h (‘Transcription factor dependent’). (E) Heat map analysis of gene expression trends across the mitochondrial UPR^mt^ transcriptome in CHOP KO, ATF4 KO, ATF5 KO and TKO cells treated with G- TPP for 12 h, calculated relative to gene expression trends of WT cells treated with G-TPP for 12 h. Enriched mitochondrial process categories identified in regulatory subgroups are labelled along the corresponding section of the heat map. (F) Compositional breakdown displayed by pie chart of the overlapping regulatory patterns of CHOP, ATF4 and ATF5 in the transcription factor dependent portion of the mitochondrial UPR^mt^ transcriptome. (G) Mitochondrial UPR^mt^ transcriptome trends in WT cells treated with G-TPP for 12 h have been separated by mitochondrial process grouping and graphed. Genes have been labelled according to the transcription factor KO lines that showed decreased gene expression relative to WT in 12 h G-TPP treated samples. Data in (A-G) represent mean data calculated from three independent experiments.

Of the 367 UPR^mt^ regulated mitochondrial genes, ∼56% were regulated by CHOP, ATF4, and ATF5 (Fig 6D), and were associated with mitochondrial processes such as OXPHOS, ROS defence, import, and translation (Figure 6E), and included mitochondrial DNA encoded genes (Figure S5D). The proportion of mitochondria associated genes regulated by one, two or all three of CHOP, ATF4, and ATF5 (Figure 6F), mirrored the observations made in the global gene analysis described above (Figure 5F). Uniquely regulated processes were also identified and included import & sorting for ATF5, and folate and pterin metabolism for ATF4 (Figure 6E). Notably, a portion of CHOP, ATF4 and ATF5 transcription factor activity involved maintaining the expression level of genes that were downregulated by the UPR^mt^ (Figure 6G). That is, genes that were downregulated by the UPR^mt^, were downregulated even further upon the loss of CHOP, ATF4 or ATF5. This occurred most strikingly for the OXPHOS gene subgroup, in which all OXPHOS genes regulated by either CHOP, ATF4, or ATF5 were further decreased in expression in transcription factor KO cells (see ‘Oxidative Phosphorylation’, Figure 6G). The three transcription factors therefore play a role in maintaining a certain level of privileged gene expression for the OXPHOS machinery that may help aid proteostasis recovery by fine tuning protein levels. For example, fine tuning of OXPHOS protein levels has been reported in *C. elegans* in which ATFS-1 downregulated OXPHOS genes (Nargund *et al*., 2015; Shpilka *et al*., 2021). However, in the mammalian system, it seems that there is a combined repression and activation of OXPHOS genes to fine tune expression. Overall, the transcriptome analyses reveal that the transcriptional program of the UPR^mt^ drives various cellular and mitochondrial processes, in which CHOP, ATF4, and ATF5 drive distinct clusters of genes that function in common processes. The analysis demonstrates that CHOP, ATF4 and ATF5 work in concert during the UPR^mt^, with the signaling arms driven by CHOP and ATF5 being related to each other, whereas ATF4’s signaling arm is quite distinct.

We noted that CHOP, ATF4 and ATF5 were dispensable for ∼44% of the global UPR^mt^ program (Figure 5E), indicating that additional UPR^mt^ transcription factors remain to be identified. The RegNetwork database was used to identify common promotor and expression patterns throughout the gene set (Liu *et al*., 2015) (Figure S5C). The analysis produced a list of 30 putative UPR^mt^ genes including MYC and MAX which have previously been shown to function in chromatin modification (Bouchard *et al*., 2001), YY1 which is known to affect mitochondrial related gene transcription (Blättler *et al*., 2012; Cunningham *et al*., 2007), and EP300 which has recently been identified to function during UPR^mt^ signaling (Li *et al*., 2021) (Figure S5C). These transcription factors represent interesting candidates for future analysis.

### The UPR^mt^ sustains PINK1/Parkin mitophagy to promote mitochondrial recovery from stress

PINK1/Parkin mitophagy and the UPR^mt^ are both activated in response to mitochondrial protein folding stress (Fiesel *et al*., 2017; Jin and Youle, 2013). To address whether there is interplay between the quality control pathways, we assessed whether PINK1/Parkin mitophagy and the UPR^mt^ influence the activity of one another. CHOP, ATF4 and ATF5 induction was analyzed by immunoblotting in response to G- TPP treatment in WT cells with and without Parkin expression (Figure 7A-H). No significant difference in CHOP, ATF4 or ATF5 levels was observed during acute proteostatic damage (Figure 7A-D), or during recovery time points (Figure 7E-H), indicating that PINK1/Parkin mitophagy does not affect UPR^mt^ signaling. Next, the influence of UPR^mt^ signaling on PINK1/Parkin mitophagy activity was assessed using the mitochondrial-targeted fluorescent reporter Keima (mtKeima) (Katayama *et al*., 2011; Lazarou *et al*., 2015; Winsor *et al*., 2020). In WT cells, PINK1/Parkin mitophagy levels peaked at 12 h G-TPP treatment and persisted at 24 h recovery before returning to baseline by 48 h recovery (Figure 7I). All UPR^mt^ deficient KO lines had slightly higher levels of mitophagy and followed a similar pattern to WT cells during the acute stress time points (4-12 h G-TPP). However, during the 24 – 48 h recovery time points, there was a large spike in mitophagy levels in CHOP, ATF4 and ATF5 KO lines (Figure 7I), demonstrating that PINK1/Parkin mitophagy is hyperactivated when the UPR^mt^ is defective. Interestingly, TKO cells did not display the same high increase in mitophagy levels as the single transcription factor KO cells (Fig 7I). Instead, only a comparatively mild elevation in mitophagy was observed during the recovery period despite TKOs having an equally or more severe proteostasis stress relative to single KOs (Figures 2-4). We assessed PINK1 accumulation (Figure 7J-K), and depolarization induced PINK1/Parkin mitophagy (Figure 7L) but did not observe any significant defects in either measure in TKO cells relative to single KOs. Receptor mediated mitophagy that is independent of PINK1/Parkin (Figure 7M), and starvation induced autophagy (Figure 7N-O), also did not show a significant defect in TKO cells relative to single KO controls, ruling out a generalized autophagy defect in the TKO line. The lowered levels of PINK1/Parkin mitophagy in TKO cells therefore appears to be a specific defect in sustaining prolonged PINK1/Parkin mitophagy activity in response to proteostasis stress (Figure 7I).

**Figure 7.**
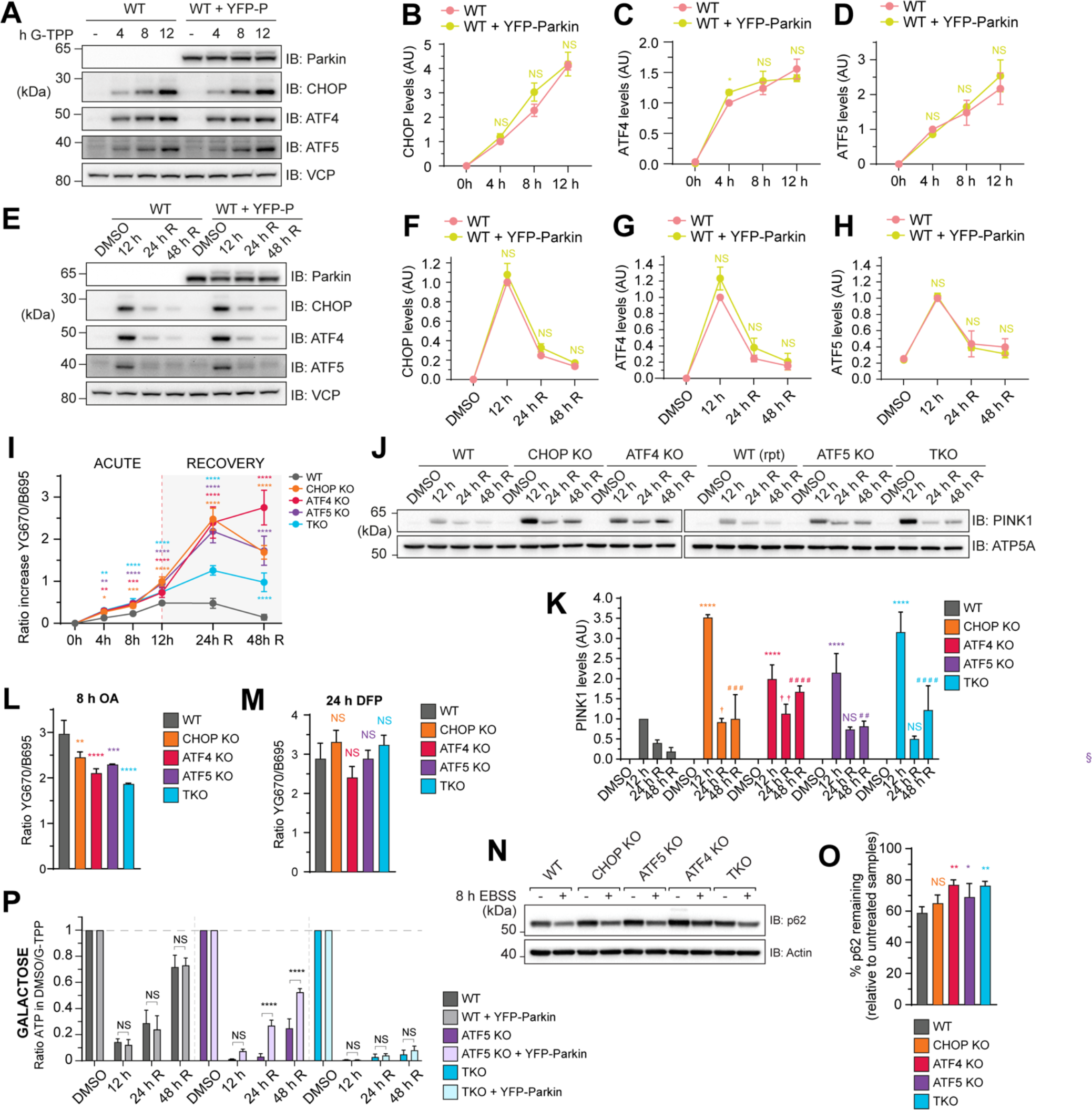
A uni-directional signaling relationship promotes PINK1/Parkin mitophagy activation during UPR^mt^-mediated recovery from proteostasis stress. (A – D) WT cells with and without YFP-Parkin (YFP-P) expression were treated with G-TPP at the indicated timepoints (4 – 12 h) and transcription factor expression was analyzed by immunoblot (IB; A) and quantified relative to WT 4 h G-TPP sample expression (B – D). (E – H) WT cells with and without YFP-Parkin expression underwent the indicated G-TPP treatment and recovery time course, and transcription factor expression was analyzed by IB (E) and quantified relative to WT 12 h G-TPP sample expression (F – H). (I) WT, CHOP KO, ATF4 KO, ATF5 KO and TKO cells expressing YFP-Parkin and mtKeima underwent the indicated acute or recovery G-TPP treatment time course and were analyzed for lysosomal-positive mtKeima using fluorescence-activated cell sorting (FACS). Ratio increase of lysosomal-positive mtKeima was calculated relative to DMSO treated samples (Winsor *et al*., 2020). (J, K) PINK1 levels in mitochondria isolated from WT, CHOP KO, ATF4 KO, ATF5 KO and TKO cells at the indicated timepoints were analyzed by IB (J) and quantified relative to WT 12 h PINK1 expression (K). (L, M) WT, CHOP KO, ATF4 KO, ATF5 KO and TKO cells expressing YFP-Parkin and mtKeima were treated with oligomycin/antimycin A (OA; L) or deferiprone (DFP; M) for the indicated times and were analyzed for lysosomal-positive mtKeima by FACS. Ratio increases of lysosomal-positive mtKeima were calculated relative to DMSO treated samples. (N, O) WT, CHOP KO, ATF4 KO, ATF5 KO and TKO cells were incubated in standard media (-) or EBSS for 8 h and levels of p62 were analyzed by IB (N) and quantified relative to standard media treated samples (O). (P) Cellular ATP levels in WT, ATF5 KO and TKO cells with and without YFP-Parkin expression in galactose-based media were analyzed at the indicated timepoints and the ratio of ATP relative to DMSO treated samples were calculated and graphed. *p<0.05, **p<0.005. ***p<0.001, ****p<0.0001 and NS = not significant ((B-D, F-H, I, P) two-way ANOVA, relative to WT; (L, M) one-way ANOVA, relative to WT; (K) two-way ANOVA, relative to WT: * = 12h, † = 24h R, # = 48h R).

To investigate whether PINK1/Parkin mitophagy functions alongside the UPR^mt^ to protect and repair mitochondrial dysfunction, OXPHOS derived cellular ATP levels were measured in WT, ATF5 KO and TKO cells with and without Parkin expression. In WT cells, PINK1/Parkin mitophagy had no effect on ATP levels across the acute stress and recovery time course (Figure 7P). In contrast, cellular ATP levels recovered significantly faster during the recovery period in ATF5 KO cells in the presence of PINK1/Parkin mitophagy (Figure 7P). TKO cells failed to recover their ATP levels even in the presence of Parkin expression (Figure 7P), consistent with their inability to drive sustained mitophagy (Figure 7I). These results demonstrate that the UPR^mt^ alone is sufficient to repair OXPHOS function during protein folding stress, but when the UPR^mt^ is defective or perhaps overwhelmed, PINK1/Parkin mitophagy responds with elevated levels that are sustained redundantly by CHOP, ATF4, and ATF5 to facilitate OXPHOS recovery. Therefore, there appears to be a unidirectional communication between the quality control pathways in which PINK1/Parkin mitophagy is influenced by the activity of the UPR^mt^ but not the other way around.

## DISCUSSION

Stressed proteomes undergo widespread remodeling that goes beyond changes in individual protein levels, including protein misfolding. Given this, it is important to capture functional changes in proteomes to better understand how cells respond to stress (Cappelletti *et al*., 2021; Maatta *et al*., 2020; Wallace *et al*., 2015; Weerapana *et al*., 2010). In MitoPQ, we have developed a functional proteomics framework that is specialized for the analysis of mitochondrial proteostasis. MitoPQ combined with knockout lines of CHOP, ATF4 and ATF5 enabled us to address some of the fundamental roles of the UPR^mt^ in maintaining mitochondrial proteostasis during protein folding stress. Our analyses show that the UPR^mt^ functions across two distinct phases of protein folding stress, beginning with a protection phase that maintains proteostasis during an insult, and then a repair phase which restores proteostasis following recovery from an insult. The role of the UPR^mt^ during the protection phase was surprising given that inhibition of protein import is thought to be a signature of mitochondrial protein folding stress (Melber and Haynes, 2018). How might the UPR^mt^ serve to protect proteostasis if the import of protective factors into mitochondria is inhibited? The answer lies in recent work revealing that changes in protein import during proteostatic stress occur temporally. It begins with a UPR^mt^-mediated boost in import early during the stress (Poveda-Huertes *et al*., 2021; Xin *et al*., 2022), followed by decreased protein import as the stress becomes more severe (Michaelis *et al*., 2022; Poveda-Huertes *et al*., 2021). Therefore, CHOP, ATF4 and ATF5 enact their UPR^mt^ program of proteostasis protection early during G-TPP treatment, prior to the more severe collapse of the import machinery that likely occurs by the end of 12 h treatment time-point. However, during recovery from the insult when G-TPP is removed, it is likely that mitochondrial protein import is re-established to enable the UPR^mt^ to function in the second phase of the UPR^mt^ involving proteostasis repair.

Mitochondria are central hubs of metabolism that generate ATP through OXPHOS. MitoPQ analyses reveal that UPR^mt^ activity is highly focused on maintaining OXPHOS metabolism during proteostasis stress (Figures 3, 4, 7). The UPR^mt^ does so directly by focusing on the protection and repair of complex I of the OXPHOS machinery, and indirectly by protecting and repairing processes that support OXPHOS metabolism, including cardiolipin biosynthesis which supports OXPHOS supercomplexes and their activity (Falabella *et al*., 2021; McKenzie *et al*., 2006), fatty acid metabolism which provides NADH, and ubiquinone biosynthesis which provides electron carriers. The mito-ribosome was also found to be highly reliant on the UPR^mt^ (Figure 3), and it is largely dedicated to the production of mtDNA encoded OXPHOS proteins, the majority of which are complex I subunits (Pagliarini *et al*., 2008). The UPR^mt^ program mediated by CHOP, ATF4 and ATF5 was additionally identified to finely tune levels of OXPHOS transcripts (Figure 6), likely to aid with syncing protein production with the protein folding capacity of mitochondria during stress. In addition to protecting and repairing OXPHOS metabolism, the UPR^mt^ through CHOP, ATF4 and ATF5 was required for elevated levels of transcripts related to one- carbon metabolism and FGF21 induction (Figure 5G, Supplementary Table 7), both of which are involved in metabolic rewiring during mitochondrial stress (Forsström *et al*., 2019). Given that deficiencies in complex I activity have been associated with Parkinson’s disease (PD) (Keeney *et al*., 2006; Schapira *et al*., 1990), it would be interesting to determine whether the UPR^mt^ contributes to preventing the progression of PD pathogenesis via maintaining OXPHOS metabolism. Indeed, there are links for the UPR^mt^ in protecting dopaminergic neurons in *C. elegans* defective in PINK1/Parkin mitophagy (Cooper *et al*., 2017), while the UPR^mt^ has also been linked to Alzheimer’s disease (Sorrentino *et al*., 2017).

It is noteworthy to highlight that proteostasis disruption of the matrix compartment resulted in a spread of the stress to the IMS when the UPR^mt^ was dysfunctional (Figure 4I-J). However, in contrast to the role of UPR^mt^ in both protecting and repairing metabolic hubs in the matrix (Figures 3,4), the UPR^mt^ was only relied upon for protection, but not repair of the IMS compartment. It is therefore likely that the IMS has its own stress response program that drives repair of the compartment (Papa and Germain, 2011).

Through MitoPQ analyses, we found that CHOP, ATF4, and ATF5 were equally important for protecting and repairing proteostasis (Figure 2, 3), leading to the question of whether they drive a singular UPR^mt^ program or whether they each govern distinct nodes of the UPR^mt^ program. Overall, the transcriptome analyses showed that CHOP, ATF4 and ATF5 act in concert with each other by driving broadly overlapping gene sets, but with each transcription factor also controlling distinct gene sets that function in common pathways. In addition, CHOP, ATF4 and ATF5 were induced independently of one another (Figure 2A-C), further supporting the conclusion that they drive independent arms of the UPR^mt^ that function in concert. The requirement for multiple transcription factors to drive the UPR^mt^ in mammals is consistent with the UPR^mt^ in *C. elegans* which is governed by ATFS-1 and DVE-1 (Nargund *et al*., 2012; Tian *et al*., 2016). However, it is likely that additional transcription factors of the mammalian UPR^mt^ remain to be identified since our transcriptome analyses revealed that CHOP, ATF4 and ATF5 drive only ∼50% of the total UPR^mt^ program (Figure 5). Multiple putative transcription factors and coregulators may be driving the remainder of the UPR^mt^ program (Figure S5C), including the histone acetyltransferase EP300 that has recently been linked to UPR^mt^ signaling in *C. elegans* and mammals (Li *et al*., 2021). In addition, large transcriptome nodes related to the regulation of histone methylation were identified in the UPR^mt^ program that were not entirely under the control of CHOP, ATF4 and ATF5 (Figure 5C), indicating that the unidentified signaling lineages may require epigenetic remodeling reminiscent of DVE-1/lin-65 mediated chromatin remodeling arm in the *C. elegans* UPR^mt^ (Tian *et al*., 2016).

PINK1/Parkin mitophagy has been observed to activate in response to proteostasis stresses that also activate the UPR^mt^ (Fiesel *et al*., 2017; Jin and Youle, 2013; Michaelis *et al*., 2022; Pimenta de Castro *et al*., 2012). However, the interplay between these pathways has been largely unclear in mammalian systems. We identified unidirectional signaling between the two quality control pathways in which PINK1/Parkin mitophagy was influenced by the UPR^mt^ but not the other way around (Figure 7). For example, PINK1/Parkin mitophagy was highly elevated in the absence of a functional UPR^mt^ (Figure 7I), but UPR^mt^ signaling was similar in the presence or absence of PINK1/Parkin mitophagy (Figure 7A-H). Our results are consistent with a model in which PINK1/Parkin mitophagy functions as a last resort quality control mechanism that disposes of mitochondria that are damaged beyond the repair capacity of the UPR^mt^. The activity of PINK1/Parkin mitophagy under conditions where the UPR^mt^ was unable to restore proteostasis is important for maintaining OXPHOS metabolism (Figure 7P), and likely also helps to prevent highly damaged mitochondria from herniating and releasing inflammatory mtDNA (McArthur *et al*., 2018; Sliter *et al*., 2018; White *et al*., 2014). Importantly, CHOP, ATF4, and ATF5, were found to redundantly sustain high levels of PINK1/Parkin mitophagy under conditions of severe proteostasis damage during the repair phase (Figure 7I), demonstrating a role for the UPR^mt^ in supporting mitophagy.

In conclusion, through the development of a functional proteomics framework in MitoPQ, we define fundamental roles for the UPR^mt^ in protecting and repairing mitochondrial proteostasis, in which OXPHOS metabolism is a key UPR^mt^ target. The transcription factors CHOP, ATF4 and ATF5 are important UPR^mt^ players that function in concert to drive ∼50% of the UPR^mt^ program that protects and repairs proteostasis, while unidirectional interplay between the UPR^mt^ and PINK1/Parkin mitophagy maintains OXPHOS activity when the UPR^mt^ is overwhelmed or dysfunctional.

## Materials and Methods

### Experimental model and subject details

All HeLa and HEK293T cell lines in this study were cultured in DMEM supplemented with 10% v/v FBS (Cytiva), 1% Penicillin-Streptomycin, 10 mM HEPES, GlutaMAX (Life Technologies) and non- essential amino acids (Life Technologies).

### Method details

#### Transfection reagents and antibodies

Transfection reagents including Lipofectamine LTX (Life Technologies) and X-tremeGENE9 (Roche) were used according to manufacturers’ instructions. All commercial antibodies used in this study are listed in the Key Resources Table.

#### Generation of knockout lines using CRISPR/Cas9 and TALEN gene editing

CHOP KO cells were generated using a transcription activator-like effector nuclease (TALEN) that targets an exon common to all splicing variants (listed Supplementary Table 1). The TALEN constructs were generated by sequential ligation of coding repeats into pcDNA3.1/Zeo-Talen(+63), as previously described (Huang *et al*., 2011). ATF4 KO, ATF5 KO, DELE1 KO and CHOP/ATF4/ATF5 TKO cells were generated using CRISPR guide RNAs (gRNAs) that target a common exon of all splicing variants of each gene. Oligonucleotides (Sigma) that contain CRISPR sequences were annealed and ligated into BbsI-linearised pSpCas9(BB)-2A-GFP vector (Ran *et al*., 2013) (a gift from Feng Zhang; Addgene plasmid # 48138). Sequence-verified gRNA constructs were then transfected into HeLa cells for 24 h and GFP-positive cells were individually sorted by fluorescence activated cell sorting (FACS) into 96 well plates. Single cell colonies were screened for the loss of the targeted gene product by immunoblotting after treatment with 300 nM Thapsigargin for 8 h (CHOP, ATF4, ATF5), or by three- primer PCR screening for genomic edits (DELE1) (Yu *et al*., 2014). The presence of frameshift indels in the genes of interest in KO clones from immunoblotting or PCR screening was confirmed by Sanger sequencing. Genomic DNA was first isolated and PCR was performed to amplify the targeted regions. For CHOP KO and ATF5 KO cells, the PCR products were subsequently cloned into a PGEM4Z vector for sequencing analysis (see Supplementary Table 2 for genotyping primers). For ATF4 KO and DELE1 KO cells, the PCR products were directly sequenced using sequencing primers that anneal to the amplified regions (Supplementary Table 2). The sequencing data for the control and the knockout cells were then analyzed using Synthego ICE v2 CRISPR Analysis Tool (https://synthego.com/products/bioinformatics/crispr-analysis) (Supplementary Table 1).

CHOP/ATF4/ATF5 TKO cells were generated by sequential transfections of CHOP TALEN plasmid into WT HeLa cells, then ATF5 CRISPR plasmid into CHOP KO cells, and then ATF4 KO CRISPR plasmid into CHOP/ATF5 DKO cells.

#### Generation of stable cell lines

pBMN-YFP-Parkin and pCHAC-mt-mKeima plasmids were described previously (Lazarou *et al*., 2015; Nguyen *et al*., 2016). Retroviruses were assembled in HEK293T cells and purified using Lentivirus Precipitation Solution (ALSTEM) as per manufacturer’s instructions. Supernatants containing purified virus were applied onto HeLa cells for 48 h in the presence of 8 μg/mL polybrene (Sigma). Following transduction, the cells were recovered in full growth media for 5 – 7 days and protein expression levels among cell lines were matched by fluorescence sorting via FACS.

#### Proteostasis stress, mitophagy and starvation treatments

For proteostasis stress experiments, cells were treated with 9 μM G-TPP (Advanced Molecular Technologies) in full growth medium for the indicated times. For proteostasis recovery experiments, after treatment with G-TPP for 12 h, cells were washed three times in excess PBS and treatment media was replaced with full growth media. Growth media was replaced with fresh growth media after 24 h recovery. For mitophagy experiments, cells were treated with 10 μM Oligomycin (Calbiochem), 4 μM Antimycin A (Sigma) and 10 μM QVD (MedChemExpress) for OA treatment, or 1 mM Deferiprone (DFP; Sigma) for DFP treatment for the indicated times. For starvation experiments, cells were fed in full media for 1 h prior to 8 h starvation in Earle’s Balanced Salt Solution (EBSS; Life Technologies).

#### Immunoblotting

Cells were lysed in 1 x LDS sample buffer (Life Technologies) in the presence of 100 mM dithiothreitol (DTT; Astral Biosciences) and heated at 99 °C with shaking for 10 min. Mitochondria were lysed in 1 x SDS sample buffer (5% w/v SDS, 10% v/v glycerol, 100 mM DTT, 50 mM Tris-Cl pH 6.8) and heated at 99 °C with shaking for 10 min. Approximately 15 – 20 μg of mitochondria or 70 μg cellular protein were subjected to NuPAGE Novex 4 – 12% Bis-Tris gels (Life Technologies) according to manufacturer’s instructions and electro-transferred to polyvinyl difluoride membranes (PVDF) prior to immunoblotting using indicated antibodies (see Supplementary Table 3 for the antibodies used in this study).

#### Mitochondrial isolation

Cell pellets were collected by scraping into cold PBS and frozen at -80 °C to increase cell lysis. Pellets were then thawed and resuspended in cold isolation solution (20 mM HEPES pH 7.6, 220 mM mannitol, 70 mM sucrose, 1 mM EDTA and 0.5 mM PMSF) and homogenized with 30 strokes in a Dounce Homogenizer. Lysates were then centrifuged at 800x g at 4 °C for 5 min to pellet nuclei and unbroken cells. Supernatants were then centrifuged at 10 000x g at 4 °C for 10 min to pellet mitochondria. Pelleted mitochondria were then washed once through resuspension in fresh isolation buffer and centrifugation at 10 000x g at 4 °C for 10 min to re-pellet mitochondria. The supernatant was removed and mitochondrial pellets were resuspended in mitochondrial storage buffer (10 mM HEPES pH 7.6, 0.5 M sucrose). Protein concentration was estimated by bicinchonic acid assay (BCA; Pierce) and aliquots of mitochondria were stored at -80 °C until use.

For oxygen consumption assays, cell pellets were collected by scraping cells into cold modified isolation buffer (70 mM sucrose, 210 mM mannitol, 1 mM EGTA, 0.5% w/v BSA (fatty acid free), 5 mM HEPES pH 7.2) and pellets were collected by centrifugation at 3 000x g for 5 min at 4 °C. Mitochondria were isolated as above with the following minor modifications: cell pellets were stored on ice prior to homogenization, and mitochondria samples were stored on ice and immediately assayed after quantification.

#### Preparation of soluble and insoluble mitochondrial protein fractions for immunoblotting

Two aliquots of 15 μg of mitochondria (‘total’ sample and ‘fractionation’ sample) were thawed on ice and pelleted by centrifugation at 10 000x g for 5 min at 4 °C. Mitochondria were then lysed on ice for 15 min in chilled lysis buffer (0.1% v/v TX-100 in PBS) at a ratio of 1 μL lysis buffer : 1 μg mitochondria. The ‘total’ fraction sample was then set aside on ice, while the ‘fractionation’ sample was centrifuged at 12 000x g for 10 min at 4 °C. Supernatants representing the soluble protein fraction were then gently removed by pipette and placed into a fresh microfuge tube that was then set aside on ice. The pelleted protein representing the insoluble protein fraction was washed through adding a volume of lysis buffer back to the microfuge tube that was equal to the volume the soluble fraction that was removed, flicking each tube gently to wash the side of the tube, centrifuging each sample at 12 000x g for 10 min at 4 °C and removing the supernatant gently by pipette. A total of two washes were performed, and after the final wash an equal volume of lysis buffer to the soluble fraction initial removed was added back to each insoluble fraction.

Each total, soluble and insoluble protein fraction for each experimental sample was warmed to room temperature and 4x SDS running buffer was added to each sample at 1x (4x SDS running buffer: 20% w/v SDS, 400 mM DTT, 40% v/v glycerol, 200 mM Tris-Cl pH 6.8). Each sample was boiled with shaking at 99 °C for 10 min, cooled to room temperature, and sonicated for 2 min in a waterbath sonicator. Samples were then subjected to NuPage Novex 4 – 12% Bis-Tris gels (Life Technologies) according to manufacturer’s instructions and electro-transferred to polyvinyl difluoride membranes (PVDF) prior to immunoblotting using indicated antibodies (Supplementary Table 3).

#### Preparation of soluble and insoluble mitochondrial protein fractions for mass spectrometry

One aliquot of 80 μg of mitochondria (fractionation sample) for each experimental sample was thawed on ice. Soluble and insoluble protein fraction isolation was performed as described earlier (see ‘Isolation of soluble and insoluble mitochondrial protein fractions for immunoblotting) up to the final addition of lysis buffer to the insoluble protein fraction. Following fractionation, 3 ng of recombinant diacylglycerol acyltransferase/mycolyltransferase (Ag85A) from Mycobacterium tuberculosis (Abcam; ab124604) per 1 μg of starting mitochondrial sample was added to each fraction in a ratiometric manner. After equilibrating to room temperature, samples were solubilized by adding 2x SDS-solubilization buffer to a final concentration of 1x (2x SDS-solubilization buffer: 10% w/v SDS, 200 mM HEPES pH 8.5). Each sample was sonicated in a waterbath sonicator for 10 min. TCEP (Pierce) to a final concertation of 10 mM and chloroacetamide (Sigma) to a final concentration of 40 mM were added to each lysate, and samples were incubated at 37 °C standing for 45 min. Lysates were then acidified by adding phosphoric acid to a final concentration of 1.2%/sample. Binding buffer (100 mM triethylammonium bicarbonate (Sigma), 90% v/v methanol, pH 7.1 with phosphoric acid (Sigma)) was added to each sample at a ratio of 1 : 7 sample volume to binding buffer. Samples were loaded onto S-Trap™ Mini columns (Protifi) and centrifuged at 6 500x g for 30 sec, repeating until the total sample was loaded with the flow-through discarded between each spin. Columns were washed by adding 400 μL binding buffer and centrifuging at 6 500x g for 30 sec. Wash steps were repeated for a total of 4 washes. Following the final wash, columns were moved to LoBind microfuge tubes (Eppendorf) and 125 μL digestion buffer (50 mM triethylammonium bicarbonate; TEAB) supplemented with sequencing grade Trypsin (Promega) at a concentration of 1 μg trypsin : 50 μg equivalent starting sample protein was added directly to each column filter. Samples were centrifuged at 1 000x g for 30 sec, and the digestion buffer flowthrough was pipetted directly back onto each column filter. Columns in their collection tubes were then sealed with parafilm and incubated statically at 37 °C for 16 hours. Digested peptides were then eluted from each column through adding to each sample 80 μL digestion buffer (no trypsin) and centrifuging samples at 3 200x g for 60 sec, adding 80 μL of 0.2% v/v formic acid (FA) and centrifuging samples at 3 200x g for 60 sec, and adding 80 μL of 50% v/v acetonitrile/0.2% v/v FA and centrifuging samples at 6 500x g for 60 sec. Each sequential eluate was pooled together within each sample, and samples were lyophilized and stored at -80 °C for downstream TMT-labelling.

#### TMT labelling and reverse-phase high pH fractionation

Lyophilized peptide pellets were reconstituted in 100 mM TEAB and concentration estimates were performed spectroscopically using a Nanodrop 1000 (Thermo Fisher Scientific). For each sample, 10 μg of peptides was aliquoted into a Lo-Bind microfuge tube (Eppendorf) and diluted to a total volume of 100 μL/sample in 100 mM TEAB. For the pooled batch control, 0.79 μg of each sample was combined in the same microfuge tube to total 100 μg peptides and diluted to 100 μL total volume in 100 mM TEAB. 10-plex TMT labels to a final quantity of 1.6 mg/label were reconstituted in acetonitrile as per manufacturer’s instruction, and 5.86 μL of the appropriate label was added to each sample according to the batch layout specified in Supplementary Table 6, while the total label volume was added to the batch control sample. Labelling reactions were performed as per manufacturer’s instructions. Labelled samples in each batch (Supplementary Table 6) were combined into a single tube, lyophilized, and stored at -80 °C. Lyophilized samples were reconstituted in 300 μL of 0.1% v/v trifluoroacetic acid (TFA) and loaded onto High pH Reversed Phase Fractionation columns (Pierce). Samples were fractionated as per manufacturer’s instructions using a modified elution gradient (Supplementary Table 4). After elution, fractions were concatenated in an equidistant manner to generate 6 sample fractions per batch sample. Concatenated samples were lyophilized and stored at -80 °C.

#### LC-MS analysis

Using a Dionex UlitMate 3000 RSLCnano system equipped with a Dionex UltiMate 3000 RS autosampler, the samples were loaded via an Acclaim PepMap 100 trap column (100μm.2cm; nanoViper; C18; 5μm; 100.; Thermo Fisher Scientific) onto an Acclaim PepMap RSLC analytical column (75μm.50cm; nanoViper; C18; 2μM; 100.; Thermo Fisher Scientific). The peptides were separated by increasing concentrations of 80% v/v ACN/0.1% v/v FA at a flow of 250 nL/min-1 for 158 min and analyzed with an Orbitrap Fusion Tribrid mass spectrometer (Thermo Fisher Scientific) operated in data-dependent acquisition mode. Each cycle was set to a fixed cycle time of 2.5 sec consisting of an Orbitrap full ms1 scan at resolution of 120 000, normalized AGC target of 50%, maximum IT of 50 ms, scan range of 380 – 1580 m/z. ms1 precursors were filtered by setting monoisotopic peak determination to peptide, intensity threshold to 5.0e3, charge state to 2 – 6 and dynamic exclusion to 60 s. Precursors were isolated in the quadrupole with a 1.6 m/z isolation window, collected to a normalized AGC target of 40% or maximum injection time (150 ms), and then fragmented with a CID collision energy of 30%. For ms3 scans, spectra were then filtered with a precursor selection range of 400 – 1200 m/z, isobaric tag loss exclusion of TMT and precursor mass exclusion set to 20 m/z low and 5 m/z high. Subsequently, 10 synchronous precursor ions were selected and scans were acquired at resolution of 50 000, normalized AGC target of 100%, maximum IT of 250 ms, and scan range of 120 – 750 m/z.

#### Calculation of mitochondrial protein solubility

Raw instrument files were processed using MaxQuant version 1.6.17 with the Andromeda search engine (Cox and Mann, 2008), searching against the Uniprot human database containing reviewed and canonical isoform variants in the FASTA format (2021), with the recombinant Ag85A sequence added as custom entry in the human database. All raw data files were analyzed using the MaxQuant proteomics data analysis workflow using the Andromeda search engine with modifications (Cox *et al*., 2011). In brief, LC-MS run was set to “Reporter ion MS” and TMT11-plex labels were set as isobaric labels with a reporter ion mass tolerance of 0.003 Da. Trypsin/P cleavage specificity was used with a maximum of 2 missed cleavages. Oxidation of methionine and N-terminal acetylation were set as variable modifications, and carbamidomethylation of cysteine was set as a fixed modification. A search tolerance of 4.5 ppm was used for ms1 and 20 ppm was used for ms2 matching. False discovery rates (FDR) were determined through the target-decoy approach and set to 1% for both peptides and proteins. The ‘Match between runs’ option was enabled, with an FDR of 1% and a mass bin size of 0.0065 Da. Minimum unique and razor peptides was set to 1, and label min. ratio count was set to 2. Data from the proteinGroups.txt output table was then normalized according to methods described previously (Plubell *et al*., 2017). The internal spiked control (Ag85A) intensities were averaged across each reporter ion channel, and this value was used to generate scaling factors for each channel to normalize reporter ion intensities for each protein to the relative starting intensities in each sample at the time of Ag85A addition. After normalization, data was imported into Perseus v1.6.15 and ‘only identified by site’, ‘reverse’ and ‘potential contaminant’ identifications were removed (Tyanova *et al*., 2016). For the TMT-labelled mass spectrometry data in Figure 1, cleaned files from Perseus were imported into R (v4.0.3) where the QRILC method of imputation was performed using the impute.LCMD package (v2.0; Lazar C, 2015) for proteins that were absent in one batch of either ‘soluble’ or ‘insoluble’ fraction groups (R Development Core Team, 2021). Cleaned data was then filtered according to a database of mitochondrial associated proteins, to only include mitochondrial proteins for downstream analysis (Kuznetsova *et al*., 2021). Computationally derived total intensities were then generated for each protein in each sample through the addition of soluble and insoluble sample fraction intensities. A percentage of total protein that was in the insoluble fraction was then calculated using the computationally derived total intensity. Aggregation index (AI) analysis was performed by calculating the average percentage of protein that is insoluble across a defined process group of mitochondrial proteins. Rate recovery analysis was performed across the 48 h recovery period for CHOP KO, ATF4 KO, ATF5 KO and TKO cells, and across the first 24 h recovery period for WT cells as WT cells had already recovered to baseline solubility by 48 h R. Solubility shifts were not calculated for proteins that were only detected in the soluble or insoluble protein fractions.

#### Mitophagy analysis using mtKeima

Cells stably expressing YFP-Parkin and mtKeima were seeded in 24 well plates 24 h prior to treatment. Treatments as indicated by the appropriate figure legends were performed as described earlier (see ‘Proteostasis stress, mitophagy and starvation treatments’). At the conclusion of the treatment time, cells were washed in 1x PBS, trypsinised and then resuspended in ice cold standard growth media. Cell pellets were centrifuged at 1 000 xg for 1.5 min and the supernatant was removed by aspiration prior to resuspension in sorting media (10% v/v FBS, 1 mM EDTA in PBS). Sample suspensions were then analyzed using the FACSDiva software on a LSR Fortessa X-20 cell sorter (BD Biosciences). Lysosomal mtKeima was measured using dual excitation ratiometric pH measurements at 488 (pH 7) and 561 (pH 4) nm lasers with 695 nm and 670 nm detection filters respectively. Additional channels used include GFP (Ex/Em: 488 nm/530 nm). Sample compensation was performed at the time of sample analysis using the FACSDiva software. A minimum of 30 000 events were collected per sample.

Data was processed using FlowJo v10.7.2. Samples were first gated to exclude debris, and then gated for single cell events using the forward and side scatter measurements. Individual event emission values for each sample were exported, and mtKeima shifts were calculated according to published methods (Winsor *et al*., 2020).

#### ATP measurements

Cells were seeded in white opaque 96 well plates (Corning) 24 h prior to treatment. Samples assayed in glucose-based media were treated with G-TPP as described earlier (see ‘Proteostasis stress treatments’).

Samples assayed in galactose-based media were placed into galactose media (Galactose media: 25 mM D-galactose, 10% v/v dialyzed FBS (Gibco), 1% v/v Penicillin-Streptomycin, 10 mM HEPES pH 7.2, GlutaMAX (Life Technologies), non-essential amino acids (Life Technologies) in glucose-free DMEM (Life Technologies)) 12 h prior to assay analysis (12 h DMSO and 12 h G-TPP) or 24 h prior to assay analysis (24 h R and 48 h R). At each analysis time point (12 h, 24 h R, 48 h R), ATP levels were assayed using the Mitochondrial ToxGlo™ Assay kit (Promega) as per manufacturer’s instructions on a FLUOstar OPTIMA (BMG LABTECH) plate reader.

#### Oxygen consumption assays

Isolated mitochondria were quantified by Bicinchoninic acid assay (Pierce) as per manufacturer’s instructions. Aliquots of 15 μg mitochondria were diluted to a final volume of 25 μL in mitochondrial assay solution (MAS; 70 mM sucrose, 220 mM mannitol, 10 mM KH2PO4, 5 mM MgCl2•6H2O, 1 mM EGTA, 0.1% (w/v) BSA (fatty acid free), 2 mM HEPES pH 7.2) and left to rest on ice. An equilibrated Seahorse cartridge plate (Agilent) was loaded with 20 mM ADP, 50 μg/μL oligomycin, 10 μM FCCP and 40 μM antimycin A, resulting in final sample plate concentrations of 2 mM, 5 μg/μL, 1 μM and 4 μM respectively. Cartridges were then incubated in a 37 °C CO2 free incubator for 45 min. During this incubation, a 25 μL aliquot of mitochondria was added to each corresponding sample well of a pre-chilled sample plate on ice. Plates were immediately centrifuged at 2 000 x g for 20 min at 4 °C and left on ice until insertion into the Seahorse XFe96 Analyzer (Agilent) after the cartridge plate calibration cycle on the analyzer was complete. Immediately prior to insertion into the Seahorse XFe96 analyzer, 155 μL of pre-warmed substrate solution (10 mM glutamate and 10 mM malate in MAS buffer) was added to the corresponding sample wells. Oxygen consumption analysis cycles were performed sequentially as follows: Basal (3 min mix, 3 min measure, 3 min mix, 3 min measure), ADP (injection, 30 sec mix, 3 min measure), oligomycin (injection, 30 sec mix, 30 sec wait, 3 min measure), FCCP (injection, 20 sec mix, 3 min measure), antimycin A (injection, 30 sec mix, 3 min measure). The protocol, from mitochondrial isolation to oxygen consumption analysis on the Seahorse analyzer, was repeated each day at the time of sample collection (12 h G-TPP (Day 1), 24 h R (Day 2), 48 h R (Day 3). Time from isolation to assay start was consistent between all three days. DMSO treated samples were analyzed alongside experimental samples at each time point. Raw data files were exported from Wave software (v2.6) and imported into Microsoft Excel. The average basal oxygen consumption of the DMSO treated samples from the first time point (12 h G-TPP) for each cell line was calculated and used to generate normalization factors to normalize variation in oxygen consumption measurements across each day of analysis. Normalized oxygen consumption rate values were then used to calculate the basal respiration rates and total respiratory capacity of each experimental sample. All data was then displayed using GraphPad Prism v9.1 and analyzed using the statistical methods listed in the corresponding figure legend.

#### qRT-PCR analysis

Total RNA was isolated using the Monarch^®^ Total RNA Miniprep Kit (New England Biolabs) as per manufacturer’s instructions. Concentrations of RNA in each sample were determined spectroscopically, and cDNA libraries were synthesized using a High-Capacity cDNA Reverse Transcription Kit (Applied Biosystems™) as per manufacturer’s instructions, using Oligo(dT)20 primers (Sigma Aldrich). Libraries were diluted 1:4 in DEPC-treated H2O. qRT-PCR analysis was performed using QuantiNova SYBR Green PCR Master Mix (Qiagen) as per manufacturer’s instructions on a RotorGene Q (Qiagen) PCR machine. Primers used in this study are located in Supplementary Table 5. Threshold values were set using Rotor-Gene Q Series Software v2.3.5. Cycle threshold (Ct) values were used to calculated fold changes in mRNA levels using the 2^-ΔΔCt^ method (Livak and Schmittgen, 2001). Transcription factor mRNA levels were normalized to GAPDH mRNA levels in each sample.

#### RNA sequencing library preparation and analysis

RNA sequencing was performed with the support of Micromon Genomics (Monash University). Total RNA samples were extracted from cell pellets using a RNeasy Mini kit (QIAGEN) as per manufacturer’s instructions. Total RNA was measured using a Qubit™ analyzer (Invitrogen™) as per manufacturer’s instructions, using 2 μL of each sample, and assayed with the Qubit™ RNA High Sensitivity Assay (Invitrogen™) as per manufacturer’s instructions. Each sample was sized and measured for RNA integrity using the Bioanalyzer 2100 (Agilent) and RNA Nano Assay kit (Agilent) as per manufacturer’s instructions. Samples were then processed using an MGI RNA Directional Library Preparation Set V2 (poly(A)), as per the manufacturer’s instructions with some modifications. Modifications were as follows: 500 ng of RNA was used as input, RNA was fragmented at 87 °C for 6 min to target an insert size of 200 – 400 bp, adapters were diluted 1/5, and the libraries were amplified with 13 cycles of PCR. The libraries were then pooled in equimolar concentrations and sequenced using an MGI DNBSEQ- 2000RS with reagent chemistry V3.1, aiming for ∼ 30 million reads per sample. Bioinformatics analysis of NGS data was performed with the support of the Monash Bioinformatics Platform (Monash University). Analysis of sequencing reads was performed using the rnasik 1.5.4 pipeline with STAR aligner, using the GRCh38 (Homo Sapiens) reference genome (Tysyganov, 2018). A UPR^mt^ induced change in gene expression was defined as changes that were statistically significant (FDR ≤ 0.05) with increased or decreased expression relative to the WT DMSO control samples of >± 0.2 fold. For mitochondrial-encoded genes, sequenced reads were trimmed with Trim Galore (--fastqc --paired -- nextera --clip_R1 1 --clip_R2 1) (https://github.com/FelixKrueger/TrimGalore) using Cutadapt 1.18 (Martin, 2011) and FastQC 0.11.9 (https://github.com/s-anders/FastQC). Trimmed reads were quantified with Salmon 1.5.2 using the selective alignment procedure (-l A --seqBias --qcBias --validateMappings) against the GENCODE vM27 transcriptome, with custom mitochondrial transcripts (properly merged bicistronic transcript sequences and corrected terminal sequences, and removal of mt-tRNAs). Transcript quantifications were summarised to gene-level with tximport (Soneson *et al*., 2015) and analysed for differential gene expression changes with DESeq2 (Love *et al*., 2014), using the apeglm (Zhu *et al*., 2018) shrinkage estimator and an lfcThreshold of log 2 (1.5). Gene expression changes with an FSOS s- value < 0.01 were considered significant. Normalized, strand-specific coverage profiles were generated with deepTools 3.5.0 bamCoverage (--filterRNAstrand [forward/reverse] --samFlagInclude 2 --samFlagExclude 256 --normalizeUsing CPM --exactScaling -bs 1 -of bigwig).

#### Gene ontology analysis

Sets of genes of interest were analyzed using the Database for Annotation, Visualization and Integrated Discovery (DAVID) online bioinformatics tool for enriched clusters of Biological Process (GOTERM_BP_DIRECT) and Molecular Function (GOTERM_MF_DIRECT) categories (Huang *et al*., 2009). The largest GO term of each cluster was used as the representative GO cluster term. Cellular pathway enrichment analysis was performed using the Kyoto Encyclopedia of Genes and Genomes (KEGG) database through the ShinyGO V0.75 bioinformatic enrichment tool (0.05 FDR cutoff) (Ge *et al*., 2019). Mitochondrial process group analysis was performed using a curated dataset of mitochondrial related proteins (Kuznetsova *et al*., 2021). Representative mitochondrial process group values were calculated by taking the average aggregation value of all detected proteins belonging to each process group in each sample.

#### Data analysis

No statistical methods were used to predetermine sample size. For western blots, band intensities were measured with ImageLab 6.1.0 (BioRad). All statistical analyses were performed using GraphPad Prism 9. Statistical significance was calculated from three independent experiments using one-way or two-way ANOVA (considered significant for p-values ≤ 0.05), as specified in the relevant Figure legend. Error bars are reported as mean ± standard deviation.

### Data availability

All methods used in this study will be submitted as protocols to https://protocols.io. The mass spectrometry proteomics data have been deposited to the ProteomeXchange Consortium via the PRIDE partner repository (Perez-Riverol et al., 2021).

## Acknowledgements

This work was supported by the National Health and Medical Research Council (NHMRC) (GNT1106471 to M.L.), the Australian Research Council (ARC) Discovery Project (DP200100347 to M.L.), and an Australian Government Research Training Program (RTP) Scholarship (to L.U.). A.F. was supported by fellowships and project grants from the NHMRC, ARC and CCWA. The study is funded by the joint efforts of The Michael J. Fox Foundation for Parkinson’s Research (MJFF) and the Aligning Science Across Parkinson’s (ASAP) initiative. MJFF administers the grant (ASAP-000350 to M.L.) on behalf of ASAP and itself.

We also thank Monash Flow Cytometry Platform (FlowCore) for sorting of cells using FACS and flow cytometry instrumentation, Micromon Genomics at Monash University for the use of NGS services and facilities, Monash Bioinformatics Platform for transcriptome analysis support, and the Monash Proteomic & Metabolomic Facility for the provision of mass spectrometry instrumentation, training and technical support. This study used BPA-enabled (Bioplatforms Australia) / NCRIS-enabled (National Collaborative Research Infrastructure Strategy) infrastructure located at the Monash Proteomics & Metabolomics Facility.

## Author contributions

L.U and M.L conceived the projects; L.U., R.B.S. and M.L designed experiments; L.U., R.L., M.S., G.K., T.N.N. performed experiments; L.U., D.L.R. and A.F. performed data analysis. L.U. and M.L. wrote the manuscript and all authors contributed to preparing and editing the manuscript.

---

**Supplementary Figure 1.**
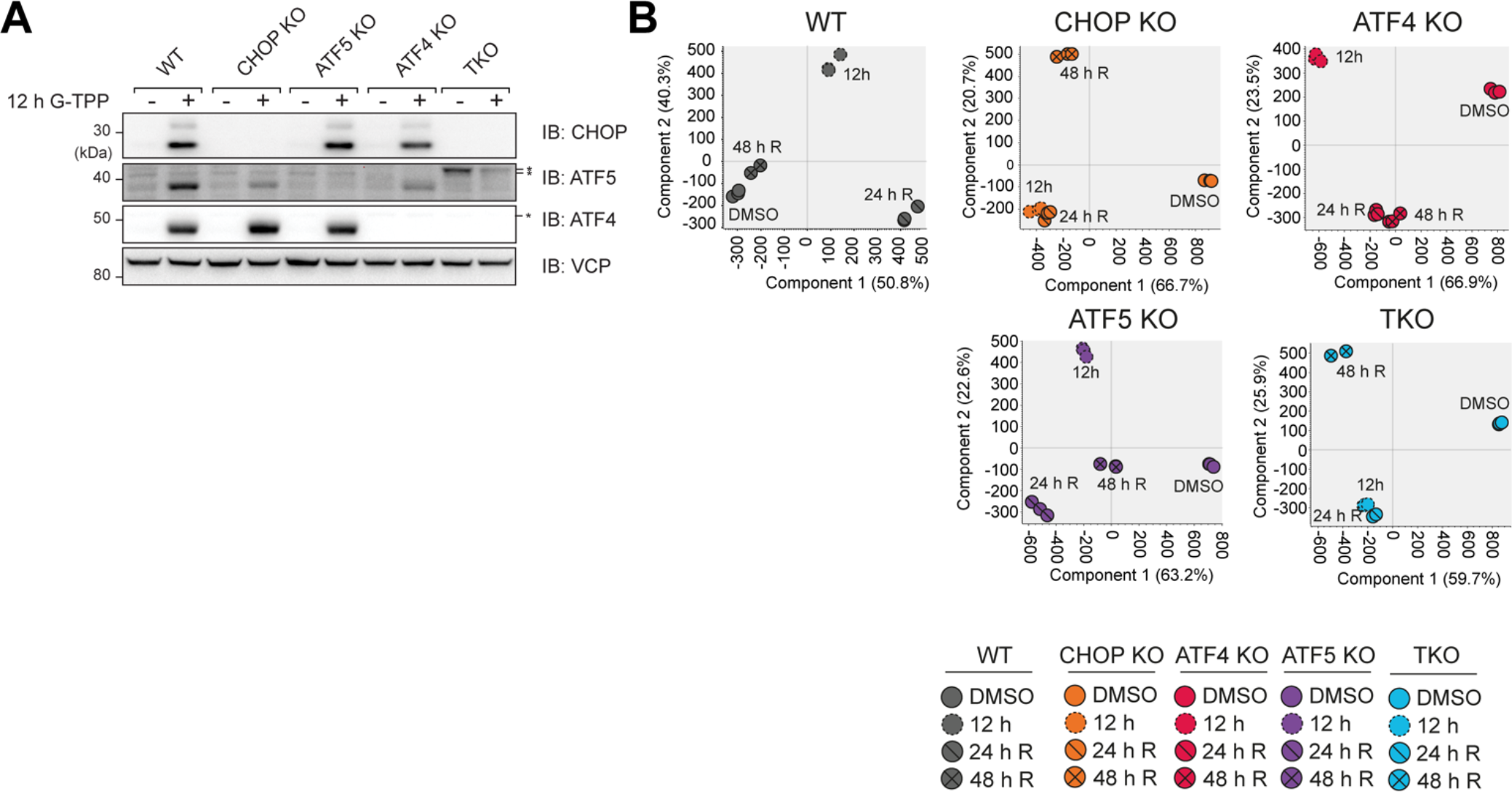
Validation of cell lines and temporal MitoPQ analysis sample trends, related to **Figure 2**. (A) WT, CHOP KO, ATF4 KO, ATF5 KO and TKO cells were treated with G-TPP for 12 h and gene expression loss in each KO cell line was validated by immunoblot. (B) Principal component analysis of MitoPQ solubility values in each temporal data set, analyzed by each cell line. Each data point in (B) represents one independent experimental sample.

**Supplementary Figure 2.**
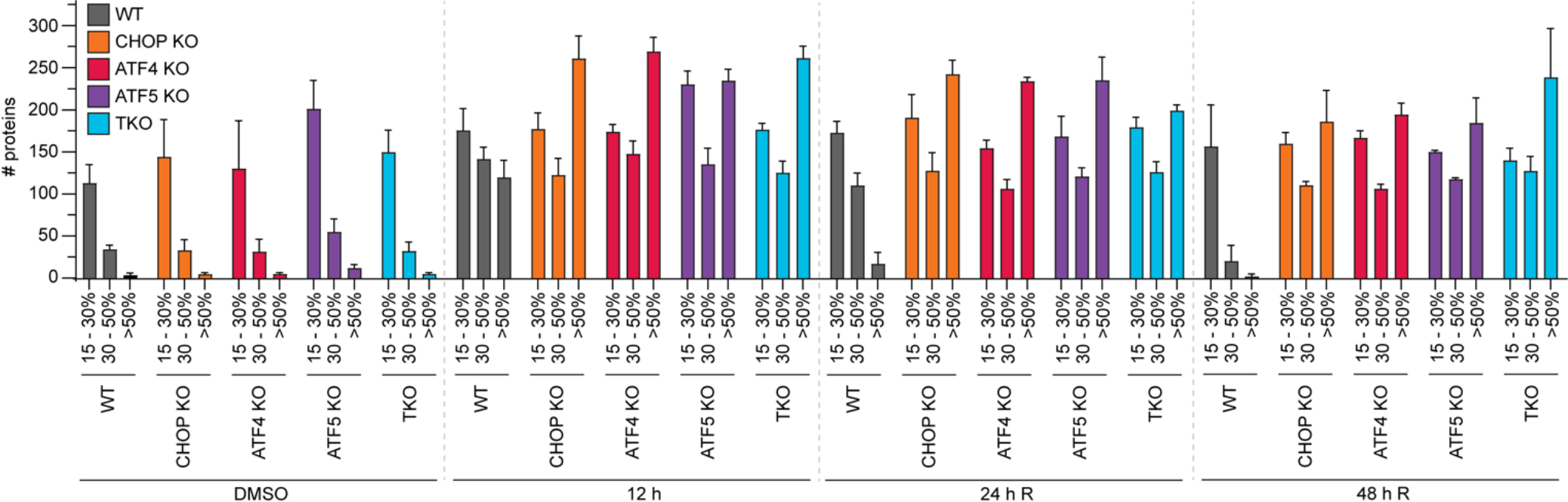
Elevated protein aggregation observed in both WT and UPR^mt^-deficient cells is largely comprised of select, strongly aggregating proteins, related to **Figures 2 and 3**. Protein aggregation trends across the mitoproteome of WT, CHOP KO, ATF4 KO, ATF5 KO and TKO cells at the indicated time points were sub-grouped into the labelled % aggregation groupings and graphed. Data represents mean data ± SD from three independent experiments.

**Supplementary Figure 3.**
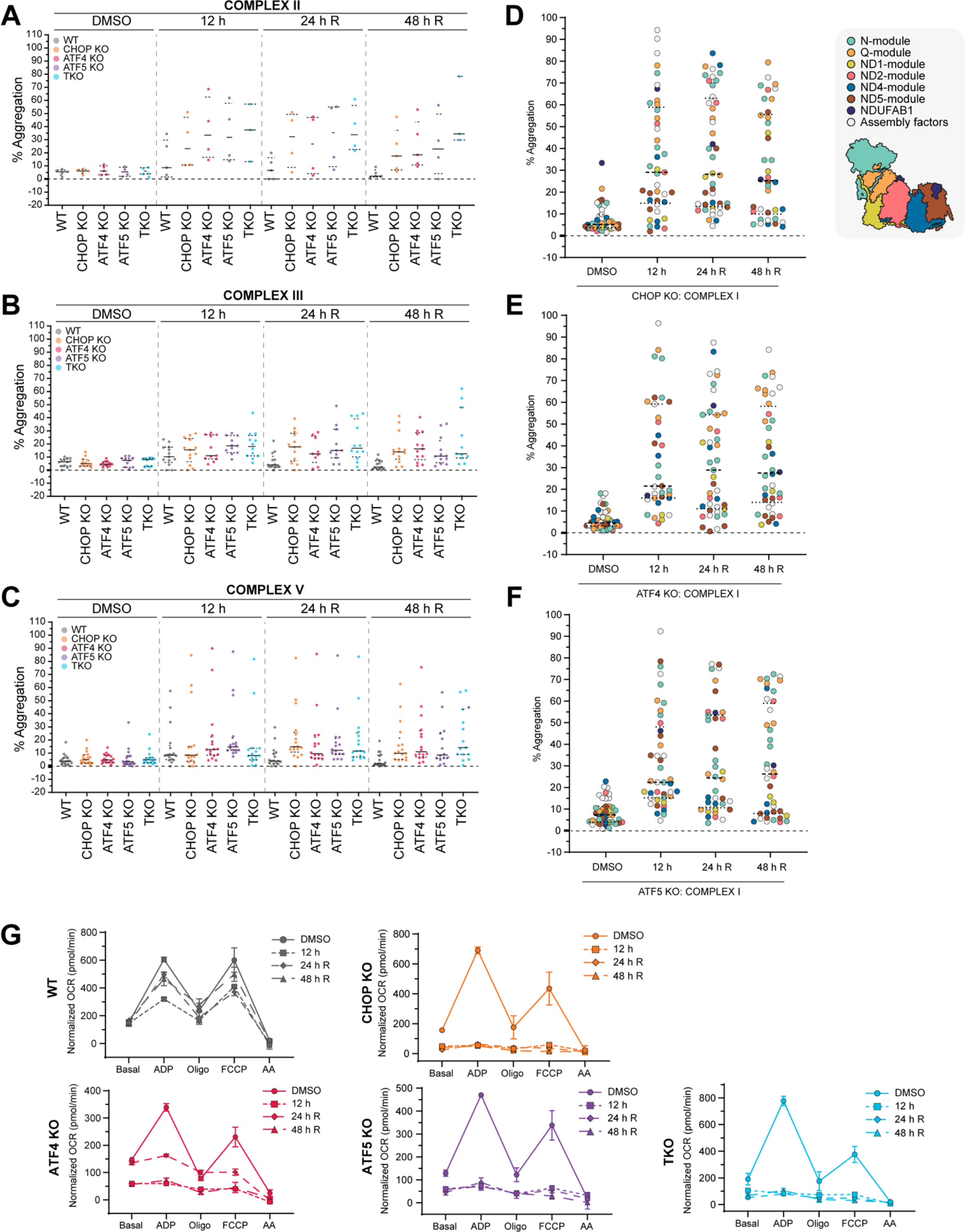
Each CHOP, ATF4 and ATF5-driven signaling arm is required for UPR^mt^-mediated OXPHOS function and protein solubility protection and repair, related to **Figure 4**. (A – C) Violin plots of the mean % aggregation of proteins comprising complex II (A), complex III (B) and complex V (C) in WT, CHOP KO, ATF4 KO, ATF5 KO and TKO cells at the indicated timepoints. (D - F) Violin plots of the mean % aggregation of proteins comprising complex I, labelled according to complex I sub-module localization in CHOP KO (D), ATF4 KO (E) and ATF5 KO (F) cells at the indicated timepoints. (G) Oxygen consumption rates (OCR; pmol/min) of mitochondria isolated from WT, CHOP KO, ATF5 KO, ATF4 KO and TKO samples used to calculate total respiratory capacity and spare respiratory capacity values in Figure 4. Data in (A – G) represent mean data ± SD (G) from three independent experiments.

**Supplementary Figure 4.**
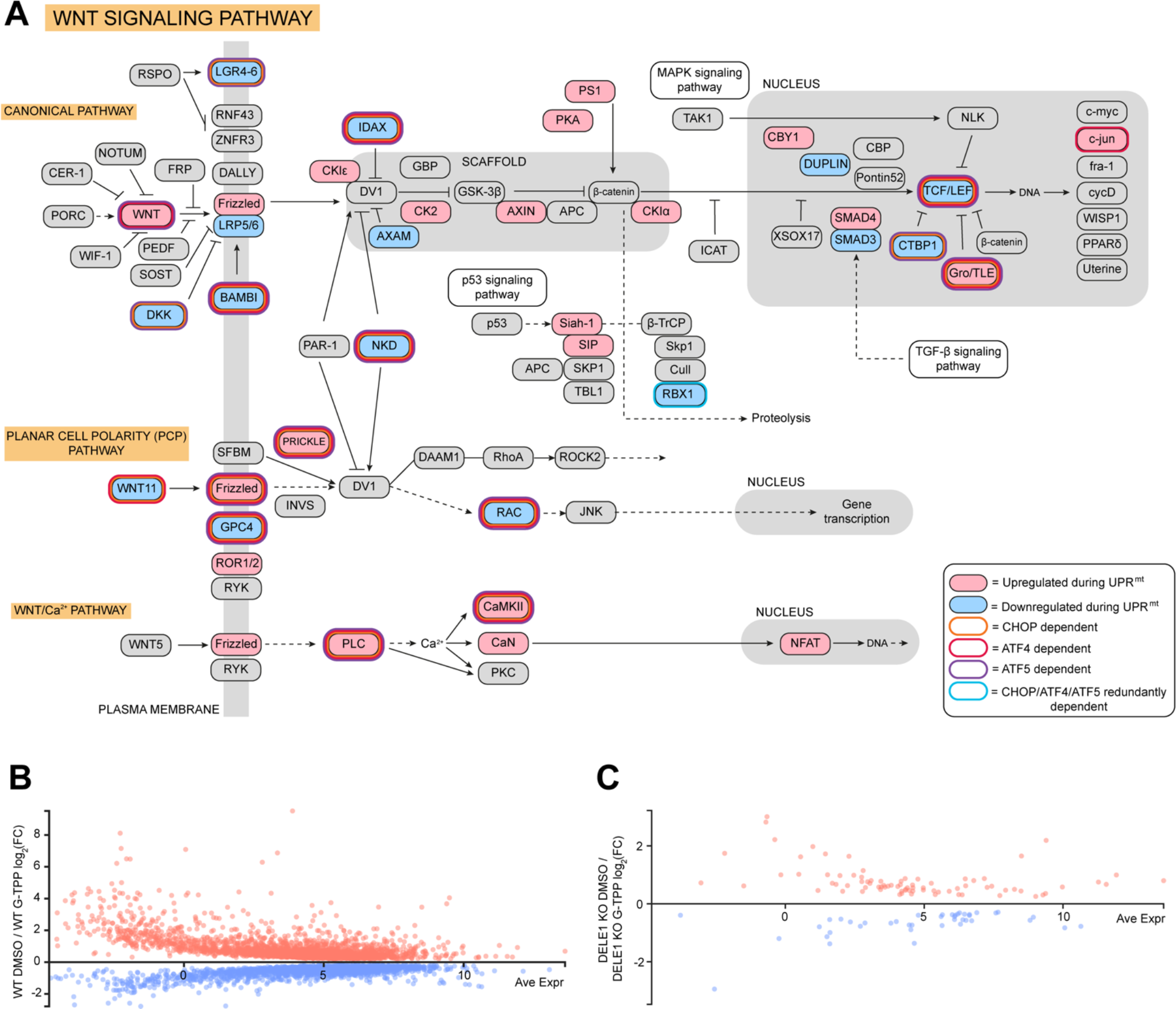
UPR^mt^ signaling regulated by CHOP, ATF4 and ATF5 drives modulation of Wnt-signaling pathway gene expression during proteostasis stress, related to **Figure 5**. (A) Wnt signaling pathway gene relationships were mapped and UPR^mt^-related expression changes including transcription factor dependency were annotated on affected genes. (B, C) Significant gene expression changes in G-TPP treated samples relative to DMSO treated samples in WT (B) and DELE1 KO (C) samples were graphed. Data in (A – C) represents mean data from three independent experiments.

**Supplementary Figure 5.**
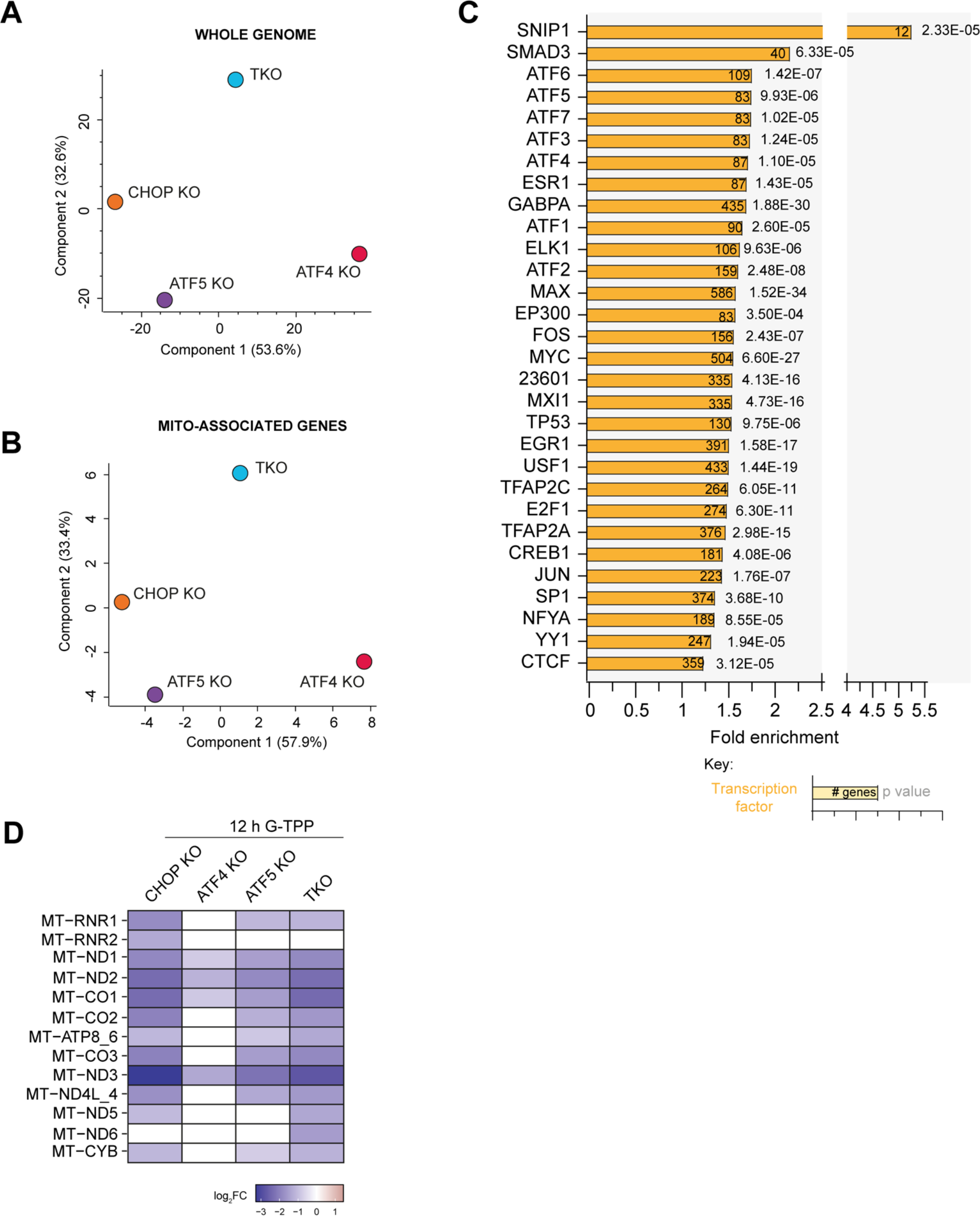
∼44% of the UPR^mt^-regulated transcriptome is under the regulatory control of unidentified signaling elements, related to **Figures 5 and 6**. (A, B) UPR^mt^ gene expression trends across the cellular genome (A) or mitochondrial genome (B) in CHOP KO, ATF4 KO, ATF5 KO and TKO cells were analyzed by Principal Component Analysis (PCA). (C) Genes that did not show reduced expression in any CHOP KO, ATF4 KO, ATF5 KO or TKO transcriptome samples were classified to be under undefined regulatory control. The gene subset with undefined regulatory control was analyzed using the RegNetwork database (Liu *et al*., 2015) through ShinyGO v0.75 (Ge *et al*., 2019) to identify transcription factors with enriched pathway and signaling representation in the undefined regulatory gene subset. (D) Changes in mtDNA-encoded rRNAs and mRNAs levels (expressed as log2 fold change (FC) of counts per million mapped) were determined by RNA-seq. The expression profiles of each gene showing statistically significant increases are in red and decreases in blue relative to WT controls for the respective treatments; non-significant changing genes are in white. Data in (A), (B) and (D) represents data from three independent experiments. Data in (C) was generated using mean data from three independent experiments.

**Supplementary Table 1.**
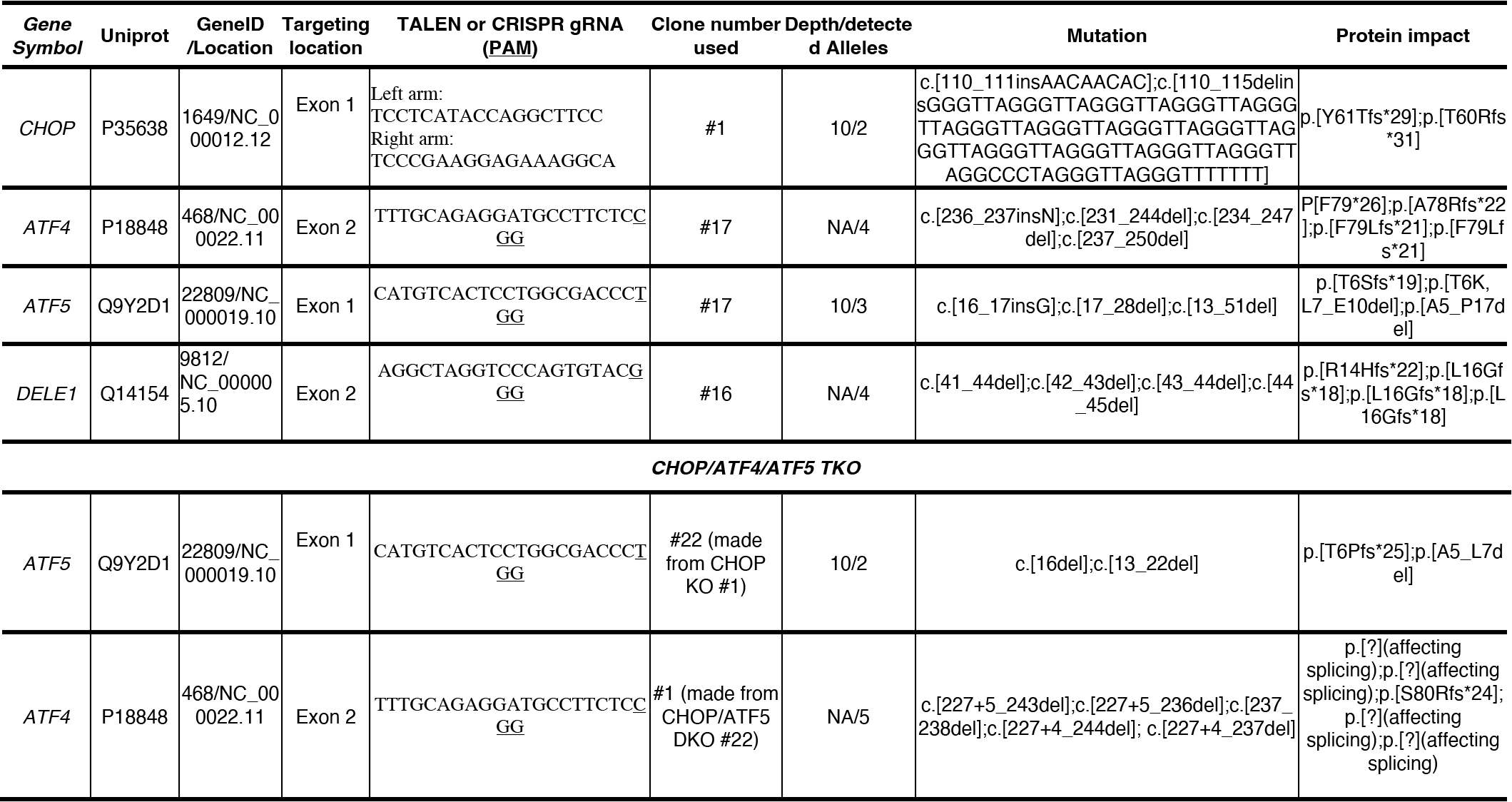
CRISPR and TALEN sequences and genotyping results of the knockout cell lines generated in this study. The indels for the targeted genes detected in the indicated knockout cell lines (“Mutation” column) and their translated proteins (“Protein impact” column) are formatted according to Human Genome Variation Society (HGVS; http://varnomen.hgvs.org/). Mutation positions within genes which have multiple splice variants are determined using the variant indicated as the canonical isoform in Uniprot. del = deletion; ins = insertion; c. = coding DNA; p. = protein; fs = frame shift; * = stop codon; [?] = affecting splicing; N= any nucleotide (A, C, T and G); X denotes one of the three amino acids A, T or S. The numbers following the asterisks denote the numbers of amino acids between the first amino acid changed after the mutation(s) and the first subsequent stop codon encountered.

**Supplementary Table 2.**
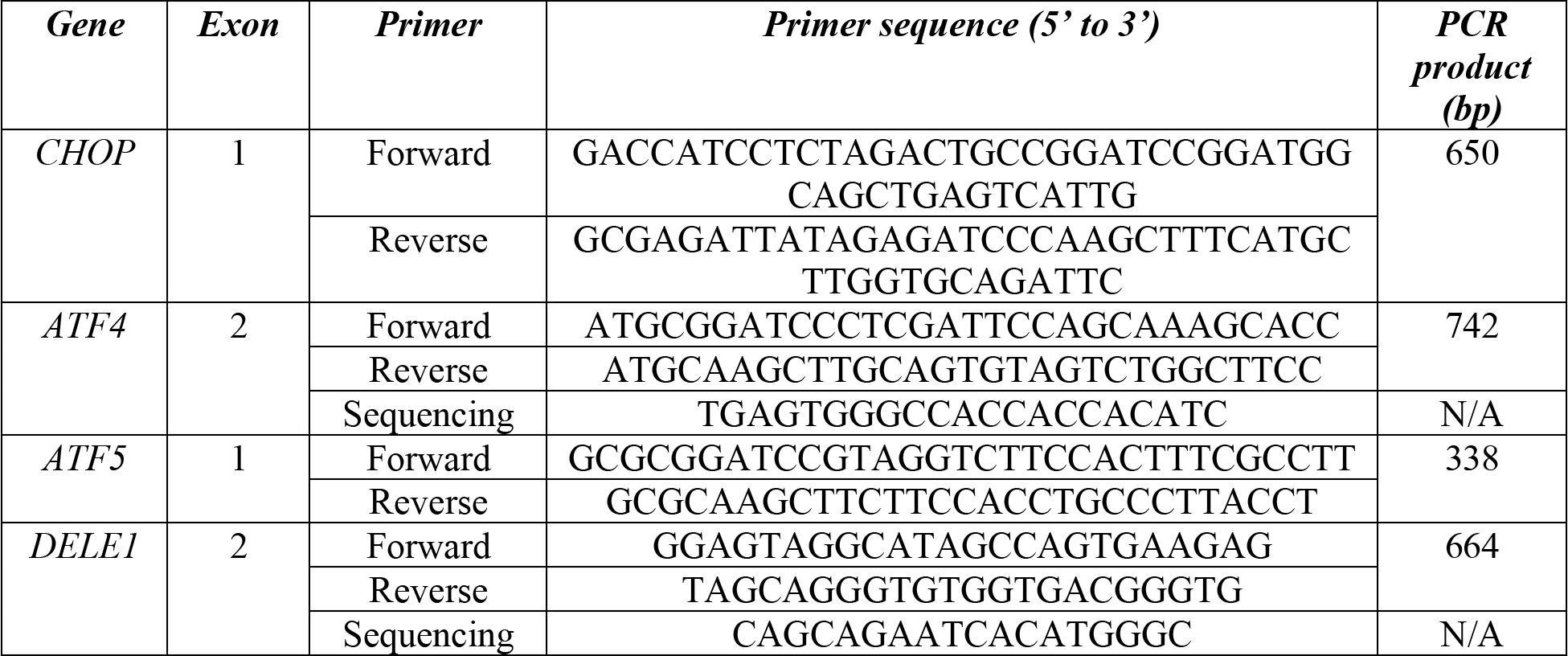
Genotyping primers for sequencing analysis of the generated knockout cell lines.

**Supplementary Table 3.** Antibodies used in this study.

| **Antibodies** | **Company** | **Species** | **Catalog #** |
| --- | --- | --- | --- |
| CHOP | Cell Signaling | Mouse | 2895 |
| ATF5 | Santa Cruz | Mouse | sc-377168 |
| ATF4 | Santa Cruz | Mouse | sc-390063 |
| PINK1 | Cell Signaling | Rabbit | 6946 |
| p62 | Abnova | Mouse | H00008878-M01 |
| Actin | Cell Signaling | Mouse | 3700 |
| PGAM5 | N/A | Rabbit | In house |
| LRPPRC | Abcam | Rabbit | ab97505 |
| AARS2 | Abcam | Rabbit | ab197367 |
| SHMT2 | Abcam | Rabbit | ab180786 |
| VDAC1 | N/A | Rabbit | In house |
| TOMM20 | Santa Cruz | Mouse | sc-17764 |
| Parkin | Santa Cruz | Mouse | sc-32282 |
| VCP | Santa Cruz | Mouse | sc-133212 |
| ATP5A | Abcam | Mouse | ab14748 |
| NDUFS1 | Proteintech | Rabbit | 12444-1-AP |
| Anti-Mouse IgG,<br>HRP-linked | Cell Signaling | Horse | 7076 |
| Anti-Rabbit IgG,<br>HRP-linked | Cell Signalling | Goat | 7074 |

**Supplementary Table 4.** Modified high-pH elution gradient used in TMT batch sample fractionation.

| **Elution step** | **% ACN** | **% TEA (0.1%)** |
| --- | --- | --- |
| Wash | 5 | 95 |
| 1 | 9 | 91 |
| 2 | 13 | 87 |
| 3 | 17 | 83 |
| 4 | 21 | 79 |
| 5 | 25 | 75 |
| 6 | 29 | 71 |
| 7 | 33 | 67 |
| 8 | 37 | 63 |
| 9 | 41 | 59 |
| 10 | 45 | 55 |
| 11 | 50 | 50 |
| 12 | 55 | 45 |

**Supplementary Table 5.**
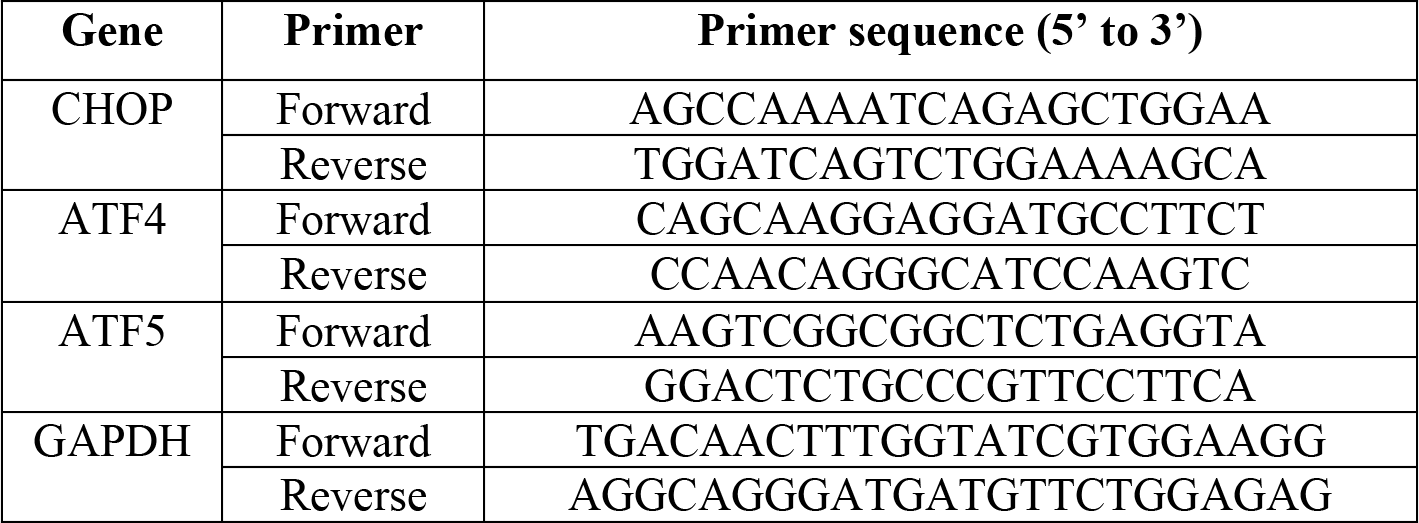
qRT-PCR primers used in this study.

**Supplementary Table 6.** TMT-batch layouts, MitoPQ data of WT 12 h G-TPP treatment samples (related to Figure 1) and WT, CHOP KO, ATF4 KO, ATF5 KO and TKO proteostasis stress and recovery samples (Related to Figures 2, 3, 4). (Provided as a separate .xlsx file)

**Supplementary Table 7.** NGS transcriptome data of WT, CHOP KO, ATF4 KO, ATF5 KO and TKO 12 h DMSO or 12 h G-TPP treated samples (related to Figures 5, 6). (Provided as a separate .xlsx file)

